# DNA from dust: comparative genomics of large DNA viruses in field surveillance samples

**DOI:** 10.1101/039479

**Authors:** Utsav Pandey, Andrew S. Bell, Daniel Renner, David A. Kennedy, Jacob Shreve, Chris L. Cairns, Matthew J. Jones, Patricia A. Dunn, Andrew F. Read, Moriah L. Szpara

## Abstract

The intensification of the poultry industry over the last sixty years facilitated the evolution of increased virulence and vaccine breaks in Marek’s disease virus (MDV-1). Full genome sequences are essential for understanding why and how this evolution occurred, but what is known about genome-wide variation in MDV comes from laboratory culture. To rectify this, we developed methods for obtaining high quality genome sequences directly from field samples without the need for sequence-based enrichment strategies prior to sequencing. We applied this to the first characterization of MDV-1 genomes from the field, without prior culture. These viruses were collected from vaccinated hosts that acquired naturally circulating field strains of MDV-1, in the absence of a disease outbreak. This reflects the current issue afflicting the poultry industry, where virulent field strains continue to circulate despite vaccination, and can remain undetected due to the lack of overt disease symptoms. We found that viral genomes from adjacent field sites had high levels of overall DNA identity, and despite strong evidence of purifying selection, had coding variations in proteins associated with virulence and manipulation of host immunity. Our methods empower ecological field surveillance, make it possible to determine the basis of viral virulence and vaccine breaks, and can be used to obtain full genomes from clinical samples of other large DNA viruses, known and unknown.

## Importance

Despite both clinical and laboratory data that show increased virulence in field isolates of MDV-1 over the last half century, we do not yet understand the genetic basis of its pathogenicity. Our knowledge of genome-wide variation between strains of this virus comes exclusively from isolates that have been cultured in the laboratory. MDV-1 isolates tend to lose virulence during repeated cycles of replication in the laboratory, raising concerns about the ability of cultured isolates to accurately reflect virus in the field. The ability to directly sequence and compare field isolates of this virus is critical to understanding the genetic basis of rising virulence in the wild. Our approaches remove the prior requirement for cell culture, and allow direct measurement of viral genomic variation within and between hosts, over time, and during adaptation to changing conditions.

## Introduction

Marek’s disease virus (MDV), a large DNA alphaherpesvirus of poultry, became increasingly virulent over the second half of the 20^th^ century, evolving from a virus that caused relatively mild disease to one that can kill unvaccinated hosts lacking maternal antibodies in as little as ten days (1–5). Today, mass immunizations with live-attenuated vaccines help to control production losses, which are mainly associated with immunosuppression and losses due to condemnation of carcasses (4, 6). Almost 9 billion broiler chickens are vaccinated against MD each year in the US alone (7). MD vaccines prevent host animals from developing disease symptoms, but do not prevent them from becoming infected, nor do they block transmission of the virus (6, 8). Perhaps because of that, those vaccines may have created conditions favoring the evolutionary emergence of the hyperpathogenic strains which dominate the poultry industry today (5). Certainly, virus evolution undermined two generations of MD vaccines (1–4). However, the genetics underlying MDV-1 evolution into more virulent forms and vaccine breaks are not well understood (4, 9). Likewise, the nature of the vaccine-break lesions that can result from human immunization with live-attenuated varicella zoster virus (VZV) vaccine is an area of active study (10–13).

Remarkably, our understanding of MDV-1 (genus: *Mardivirus,* species: Gallid alphaherpesvirus type 2) genomics and genetic variation comes exclusively from the study of 10 different laboratory-grown strains (14–21). Most herpesviruses share this limitation, where the large genome size and the need for high-titer samples has led to a preponderance of genome studies on cultured virus, rather than clinical or field samples (22–28). Repeated observations about the loss of virulence during serial passage of MDV-1 and other herpesviruses raises concerns about the ability of cultured strains to accurately reflect the genetic basis of virulence in wild populations of virus (25, 29–31). The ability to capture and sequence viral genomes directly from host infections and sites of transmission is the necessary first step to reveal when and where variations associated with vaccine-breaks arise, which one(s) spread into future host generations, and to begin to understand the evolutionary genetics of virulence and vaccine failure.

Recent high-throughput sequencing (HTS) applications have demonstrated that herpesvirus genomes can be captured from human clinical samples using genome amplification techniques such as oligonucleotide enrichment and PCR amplicon-based approaches (32–36). Here we present a method for the enrichment and isolation of viral genomes from dust and feather follicles, without the use of either of these solution-based enrichment methods. Chickens become infected with MDV by the inhalation of dust contaminated with virus shed from the feather follicles of infected birds. Although these vaccinated hosts were infected by and shedding wild MDV-1, there were no overt disease outbreaks. Deep sequencing of viral DNA from dust and feather follicles enabled us to observe, for the first time, the complete genome of MDV-1 directly from field samples of naturally infected hosts. This revealed variations in both new and known candidates for virulence and modulation of host immunity. These variations were detected both within and between the virus populations at different field sites, and during sequential sampling. One of the new loci potentially associated with virulence, in the viral transactivator ICP4 (MDV084 / MDV100), was tracked using targeted gene surveillance of longitudinal field samples. These findings confirm the genetic flexibility of this large DNA virus in a field setting, and demonstrate how a new combination of HTS and targeted Sanger-based surveillance approaches can be combined to understand viral evolution in the field.

## Results

### Sequencing, assembly and annotation of new MDV-1 consensus genomes from the field

To assess the level of genomic diversity within and between field sites that are under real world selection, two commercial farms in central Pennsylvania (11 miles apart) with a high prevalence of MDV-1 were chosen **(Figure 1A).** These operations raise poultry for meat (also known as broilers), and house 25,000-30,000 individuals per house. The poultry were vaccinated with a bivalent vaccine composed of MDV-2 (strain SB-1) and HVT (strain FC126). In contrast to the Rispens vaccine, which is an attenuated MDV-1 strain, MDV-2 and HVT can be readily distinguished from MDV-1 across the length of the genome, which allowed us to differentiate wild MDV-1 from concomitant shedding of vaccine strains. These farms are part of a longitudinal study of MDV-1 epidemiology and evolution in modern agricultural settings (37, 38).

**Figure 1.**
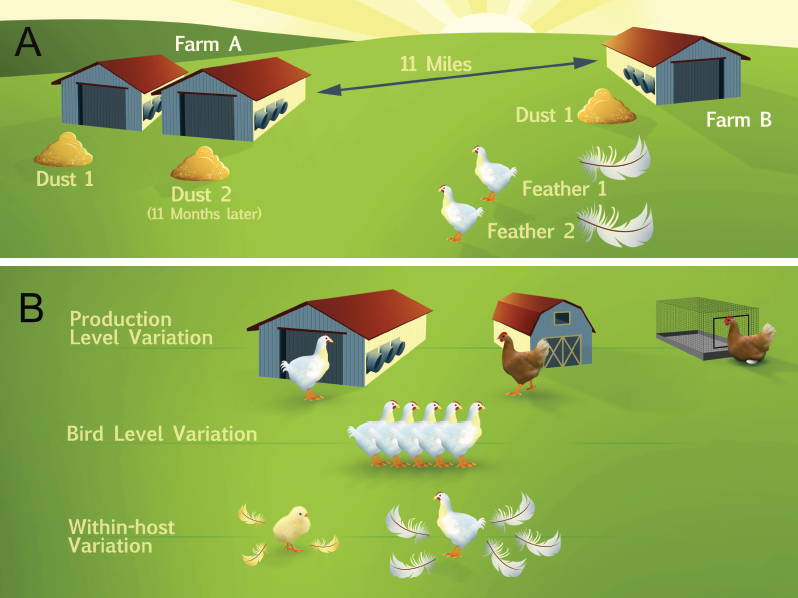
Diagram of samples collected for genome sequencing of field isolates of MDV. **(A)** Samples collected for genome sequencing were sourced from two Pennsylvania farms with large-scale operations that house approximately 25,000-30,000 individuals per building. These farms were separated by 11 miles. On Farm A, two separate collections of dust were made 11 months apart. On Farm B, we collected one dust sample and individual feathers from several hosts, all at a single point in time. In total, three dust collections and two feathers were used to generate five consensus genomes of MDV field isolates **(Table 1)**. **(B)** These methods can be used to explore additional aspects of variation in future studies. (Artwork by Nick Sloff, Penn State University, Department of Entomology).

To obtain material for genomic surveillance, we isolated MDV nucleocapsids from dust or epithelial tissues from the individual feather follicles of selected hosts (see Methods and **Supplemental Figure S1–S2** for overviews, and **Supplemental Tables S1–S2** for DNA yields). A total of five uncultured wild-type samples of MDV were sequenced using an in-house Illumina MiSeq sequencer **(Table 1,** lines 4-6; see **Methods** for details). The sequence read data derived from dust contained approximately 2-5% MDV-1 DNA, while the feather samples ranged from ~27%-48% MDV-1 **(Table 1,** line 6). Since dust represents the infectious material that transmits MDV from host to host, and across generations of animals that pass through a farm or field site, we pursued analysis of wild MDV-1 genomes from both types of source material.

**Table 1:** Field sample statistics and assembly of MDV-1 consensus genomes.

| Line # | Category<sup>a</sup> | Farm A -<br>dust 1 | Farm A -<br>dust 2 | Farm B -<br>dust | Farm B -<br>feather 1 | Farm B -<br>feather 2 |
| --- | --- | --- | --- | --- | --- | --- |
| 1 | **Nanograms of DNA** | 120 | 127 | 144 | 12 | 27 |
| 2 | **% MDV-1** | 2.4% | 1.3% | 0.6% | 40.6% | 5.7% |
| 3 | **% MDV-2** | 4.6% | 2.7% | 5.9% | 0.1% | 0% |
| 4 | **Total Reads<sup>b</sup>** | $1.4 \times 10^7$ | $2.5 \times 10^7$ | $2.7 \times 10^7$ | $3.9 \times 10^5$ | $3.4 \times 10^5$ |
| 5 | **MDV-specific reads<sup>b</sup>** | $3.7 \times 10^5$ | $5.1 \times 10^5$ | $1.4 \times 10^6$ | $1.0 \times 10^5$ | $1.7 \times 10^5$ |
| 6 | **% MDV specific reads** | 2.6% | 2.0% | 5.2% | 26.9% | 48.3% |
| 7 | **Average depth (X-fold)** | 271 | 333 | 597 | 44 | 68 |
| 8 | **Genome length** | 177,967 | 178,049 | 178,169 | 178,327 | 178,540 |
| 9 | **NCBI accession**<br>**number** | KU173116 | KU173115 | KU173119 | KU173117 | KU173118 |

<sup>a</sup>Lines 1-3 refer to sample preparation, lines 4-6 to Illumina MiSeq output, and lines 7-9 to new viral genomes.
<sup>b</sup>Sequence read counts in line 4 and 5 are the sum of forward and reverse reads for each sample.

Consensus genomes were created for each of the five samples in **Table 1,** using a recently described combination of *de novo* assembly and reference-guided alignment of large sequence blocks, or contigs **(Figure 2A)** (39). Nearly complete genomes were obtained for all five samples **(Table 1).** The coverage depth for each genome was directly proportional to the number of MDV-1 specific reads obtained from each sequencing library **(Table 1,** line 5,7). The dust sample from Farm B had the highest coverage depth, at an average of almost 600X across the viral genome. Feather 1 from Farm B had the lowest coverage depth, averaging 44X genome-wide, which still exceeds that of most bacterial or eukaryotic genome assemblies. The genome length for all 5 samples was approximately 180 kilobases **(Table 1),** which is comparable to all previously sequenced MDV-1 isolates (14–21).

**Figure 2.**
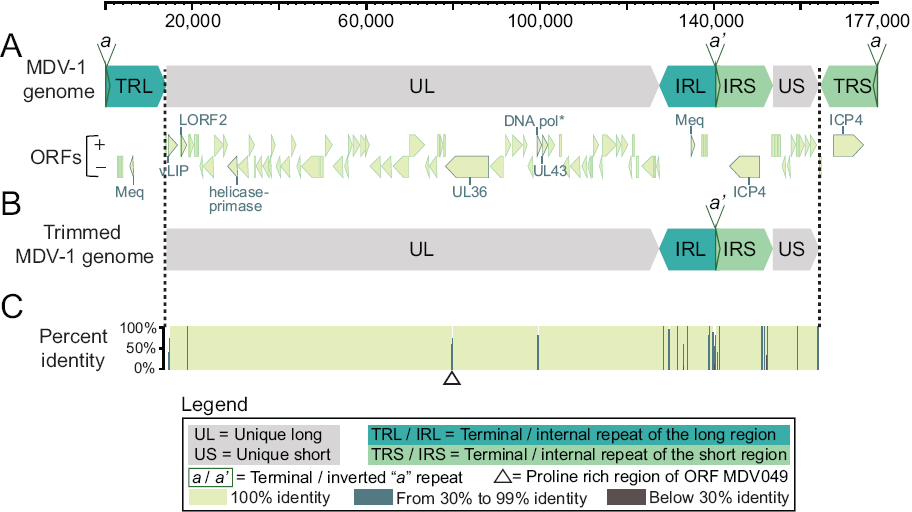
The complete MDV-1 genome includes two unique regions and two sets of large inverted repeats. **(A)** The full structure of the MDV-1 genome includes a unique long region (UL) and a unique short regions (US), each of which are flanked large repeats known as the terminal and internal repeats of the long region (TRL and IRL) and the short region (TRS and IRS). Most ORFs (pale green arrows) are located in the unique regions of the genome. ORFs implicated in MDV pathogenesis are outlined and labeled; these include ICP4 (MDV084 / MDV100), UL36 (MDV049), and Meq (MDV005 / MDV076) (see Results for complete list). **(B)** A trimmed genome format without the terminal repeat regions was used for analyses in order to not over-represent the repeat regions. **(C)** Percent identity from mean pairwise comparison of five consensus genomes, plotted spatially along the length of the genome. Darker colors indicate lower percent identity (see Legend).

For each field sample collected and analyzed here, we assembled a consensus viral genome. We anticipated that the viral DNA present in a single feather follicle might be homotypic, based on similar results found for individual vesicular lesions of the alphaherpesvirus VZV (10, 33). We further expected that the genomes assembled from a dust sample would represent a mix of viral genomes, summed over time and space. Viral genomes assembled from dust represent the most common genome sequence, or alleles therein, from all of the circulating MDV-1 on a particular farm. The comparison of consensus genomes provided a view into the amount of sequence variation between Farm A and Farm B, or between two individuals on the same Farm **(Table 2)**. In contrast, examining the polymorphic loci within each consensus genome assembly allowed us to observe the level of variation within the viral population at each point source **(Figures 3–4, Supplemental Figure S3, Supplemental Table S3**).

**Figure 3:**
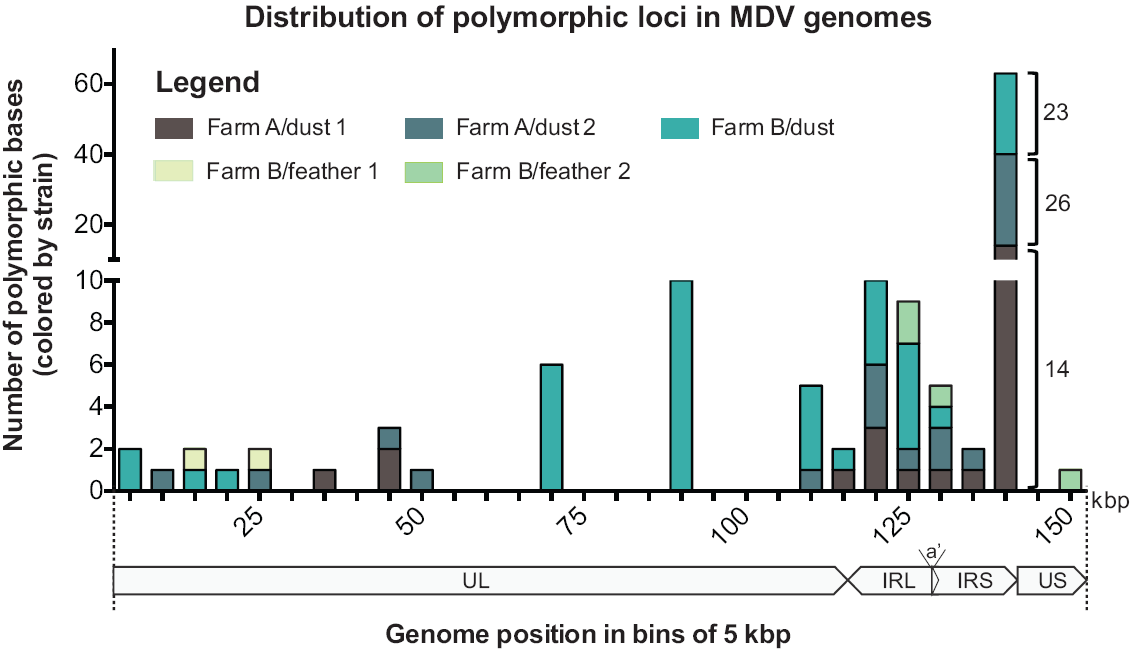
Genome-wide distribution of polymorphic bases within each consensus genome. Polymorphic base calls from each MDV genome were grouped in bins of 5 kb and the sum of polymorphisms in each bin was plotted. Farm B-dust (aqua) contained the largest number of polymorphic bases, with the majority occurring in the repeat region (IRL/IRS). Farm A-dust 1 (brown) and Farm A-dust 2 (gray) harbored fewer polymorphic bases, with similar distribution to Farm B-dust. Polymorphic bases detected in feather genomes were more rare, although this likely reflects their lower coverage depth (see **Table 1**). Note that the upper and lower segments of the y-axis have different scales; the number of polymorphic bases per genome for the split column on the right are labeled for clarity.

**Figure 4.**
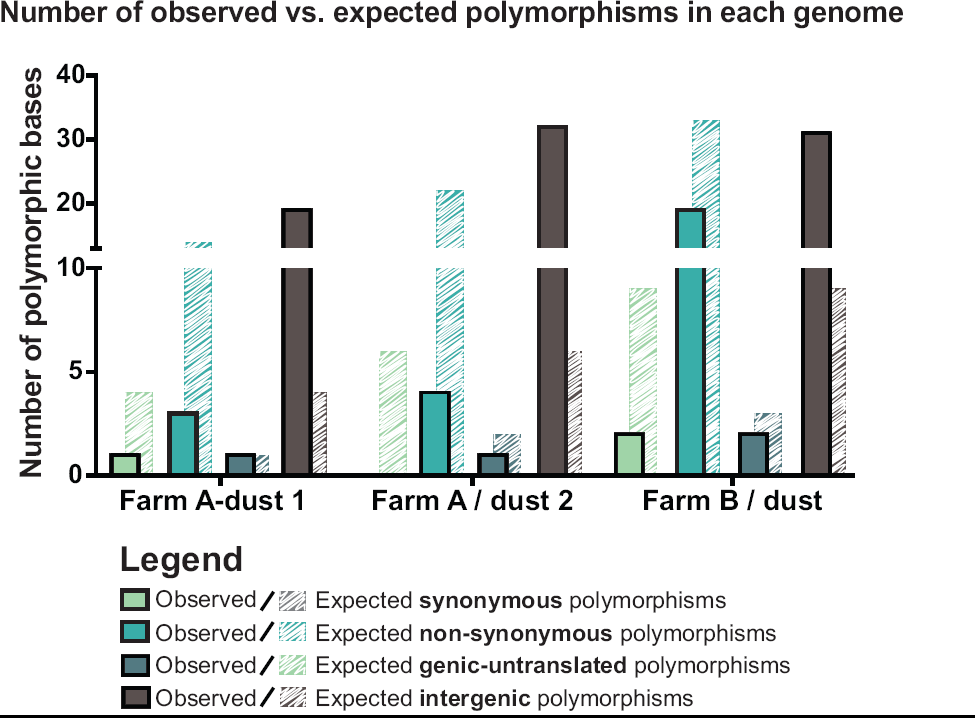
Observed vs. expected polymorphism categories for each consensus genome. Each consensus genome was analyzed for the presence of polymorphic loci (see Methods for details). Observed polymorphic loci (solid bars) were categorized as causing synonymous (green) or non synonymous (aqua) mutations, or as genic-untranslated (gray) or intergenic (brown). The expected outcomes (striped bars) for a random distribution of polymorphisms is plotted behind the observed outcomes (solid bars) for each category. For all genomes there was a significant difference of the observed-vs.-expected intergenic polymorphisms, relative to those of other categories.

### DNA and amino acid variations between five new field genomes of MDV-1

We began our assessment of genetic diversity by determining the extent of DNA and amino acid variations between the five different consensus genomes. We found that the five genomes are highly similar to one another at the DNA level, with the percent homology ranging from 99.4% to 99.9% in pairwise comparisons **(Figure 2C, Table 2).** These comparisons used a trimmed genome format **(Figure 2B)** where the terminal repeat regions had been removed, so that these sequences were not over-represented in the analyses. The level of identity between samples is akin to that observed in closely related isolates of herpes simplex virus 1 (HSV-1) (39). Observed nucleotide differences were categorized as genic or intergenic, and further sub-divided based on whether the differences were insertions or deletions (INDELs) or single-nucleotide polymorphism (SNPs) **(Table 2).** The number of nucleotide differences was higher in intergenic regions than in genic regions for all genomes. For the INDEL differences, we also calculated the minimum number of events that could have led to the observed differences, to provide context on the relative frequency of these loci in each genome. We anticipate that these variations include silent mutations, as well as potentially advantageous or deleterious coding differences.

**Table 2.** Pair-wise DNA identity and variant proteins between pairs of consensus genomes.

| Comparisons | % DNA identity | Total # bp different | Intergenic |  | Genic |  |  |
| --- | --- | --- | --- | --- | --- | --- | --- |
|  |  |  | INDELs (# events) | SNPs | INDELs (# events) | Synonymous SNPs | Non-synonymous SNPs |
| Different farms: Dust vs. dust |  |  |  |  |  |  |  |
| Farm B-dust vs. Farm A-dust 1 | 99.73 | 353 | 143 (22) | 140 | 66 (1) in DNA-pol<sup>a</sup> | 1 in helicase-primase<sup>a</sup> | 3 (one each in vLIP, LORF2, UL43)<sup>a</sup> |
| Farm B-dust vs. Farm A-dust 2 | 99.87 | 195 | 49 (14) | 76 | 66 (1) in DNA-pol<sup>a</sup> | 1 in helicase-primase<sup>a</sup> | 3 (one each in vLIP, LORF2, UL43)<sup>a</sup> |
| Same farm, same time: Dust vs. host |  |  |  |  |  |  |  |
| Farm B-dust vs. Farm B-feather 1 | 99.64 | 552 | 476 (11) | 6 | 66 (1) in DNA-pol<sup>a</sup> | 1 in helicase-primase<sup>a</sup> | 3 (one each in vLIP, LORF2, UL43)<sup>a</sup> |
| Farm B-dust vs. Farm B-feather 2 | 99.52 | 687 | 572 (19) | 45 | 66 (1) in DNA-pol<sup>a</sup> | 1 in helicase-primase<sup>a</sup> | 3 (one each in vLIP, LORF2, UL43)<sup>a</sup> |
| Same farm: Separated in time and space |  |  |  |  |  |  |  |
| Farm A-dust 1 vs. Farm A-dust 2 | 99.76 | 338 | 170 (20) | 168 | 0 | 0 | 0 |
| Same farm, same time: one host vs. another |  |  |  |  |  |  |  |
| Farm B-feather 1 vs. Farm B-feather 2 | 99.38 | 973 | 972 (9) | 1 | 0 | 0 | 0 |

<sup>a</sup> Abbreviations refer to DNA polymerase processivity subunit protein UL42 (MDV055); helicase-primase subunit UL8 (MDV020); vLIP, lipase homolog (MDV010); LORF2, immune evasion protein (MDV012); UL43 membrane protein (MDV056)

To understand the effect(s) of these nucleotide variations on protein coding and function, we next compared the amino acid (AA) sequences of all open reading frames (ORFs) for the five isolates. The consensus protein coding sequences of all five isolates were nearly identical, with just a few differences **(Table 2).** In comparison to the other four samples, Farm B-dust harbored AA substitutions in four proteins. A single non-synonymous mutation was seen in each of the following: the virulence-associated lipase homolog vLIP (MDV010; Farm B-dust, S501 A) (40), the MHC class I immune evasion protein LORF2 (MDV012; Farm B-dust, L311W) (41), and the probable membrane protein UL43 (MDV056; Farm B-dust, S74L). A single synonymous mutation was observed in the DNA helicase-primase protein UL8 (MDV020; Farm B-dust, L253L). Finally, a 22 AA insertion unique to Farm B-dust was observed in the DNA polymerase processivity subunit protein UL42 (MDV055; Farm B-dust, insertion at AA 277). We did not observe any coding differences between temporally separated dust isolates from Farm A or between feather isolates from different hosts in Farm B, although both of these comparisons **(Table 2,** bottom) revealed hundreds of noncoding differences.

### Detection of polymorphic bases within each genome

Comparing viral genomes found in different sites provides a macro-level assessment of viral diversity. We next investigated the presence of polymorphic viral populations within each consensus genome, to reveal how much diversity might exist within a field site (as reflected in dust-derived genomes) or within a single host (as reflected in feather genomes).

For each consensus genome, we used polymorphism detection analysis to examine the depth and content of the sequence reads at every nucleotide position in each genome (see **Methods** for details). Rather than detecting differences between isolates, as in **Table 2,** this approach revealed polymorphic sites within the viral population that contributed to each consensus genome. We detected 2-58 polymorphic sites within each consensus genome **(Figure 3)** (see **Methods** for details). The feather genomes had a lower number of polymorphisms compared to the dust genomes, which may be due to low within-follicle diversity or the relatively low sequence coverage. INDELs were not included in this polymorphism analysis, but clearly contributed to between-sample variation **(Table 2),** suggesting that this may be an underestimate of the overall amount of within-sample variation. Viral polymorphisms were distributed across the entire length of the genome **(Figure 3),** with the majority concentrated in the repeat regions. Application of a more stringent set of parameters (see **Methods** for details) yielded a similar distribution of polymorphisms, albeit with no polymorphisms detected in feather samples due to their lower depth of coverage **(Supplemental Figure S3).** These data reveal that polymorphic alleles are present in field isolates, including in viral genomes collected from single sites of shedding in infected animals.

To address the potential effect(s) of these polymorphisms on MDV biology, we divided the observed polymorphisms into categories of synonymous, non-synonymous, genic-untranslated, or intergenic **(Supplemental Table S3).** The majority of all polymorphisms were located in intergenic regions **(Supplemental Table S3).** We next investigated whether evidence of selection could be detected from the distribution of polymorphisms in our samples. One way to assess this is to determine whether the relative frequencies of synonymous, non-synonymous, genic-untranslated, and intergenic polymorphisms can be explained by random chance. If the observed frequencies differ from those expected from a random distribution, it would suggest genetic selection. After calculating the expected distribution in each sample (as described in **Methods),** we determined that the distribution of variants differed from that expected by chance in each of our dust samples **(Figure 4,** Farm A-Dust 1: λ^2=^68.16, d.f. =3, p<0.001; Farm A-Dust 2: χ^2^=128.57, d.f. =3, p<0.001; Farm B-Dust **1:** λ^2^=63.42, d.f. =3, p<0.001). In addition, we found in pairwise tests that the number of observed intergenic polymorphisms was significantly higher than the observed values for other categories **(Supplemental Table S4).** This suggests that the mutations that occurred in the intergenic regions were better tolerated and more likely to be maintained in the genome; *i.e.* that purifying selection was acting on coding regions.

### Tracking shifts in polymorphic loci over time

In addition to observing polymorphic SNPs in each sample at a single moment in time, we explored whether any shifts in polymorphic allele frequency were detected in the two sequential dust samples from Farm A. We found one locus in the ICP4 (MDV084 / MDV100) gene (nucleotide position 5,495) that was polymorphic in the Farm A-dust 2 sample, with nearly equal proportions of sequence reads supporting the major allele (C) and the minor allele (A) **(Figure 5A).** In contrast, this locus had been 99% A and only 1% C in Farm A-dust 1 (collected 11 months earlier in another house on the same farm), such that it was not counted as polymorphic in that sample by our parameters (see **Methods** for details). At this polymorphic locus, the nucleotide C encodes a serine, while nucleotide A encodes a tyrosine. The encoded AA lies in the C-terminal domain of ICP4 (AA position 1,832). ICP4 is an important immediate-early protein in all herpesviruses, where it serves as a major regulator of viral transcription (42–44). The role of ICP4 in MDV pathogenesis is also considered crucial because of its proximity to the latency associated transcripts (LAT) and recently described miRNAs (44–46). In a previous study of MDV-1 attenuation through serial passage *in vitro*, mutations in ICP4 appeared to coincide with attenuation (31).

**Figure 5:**
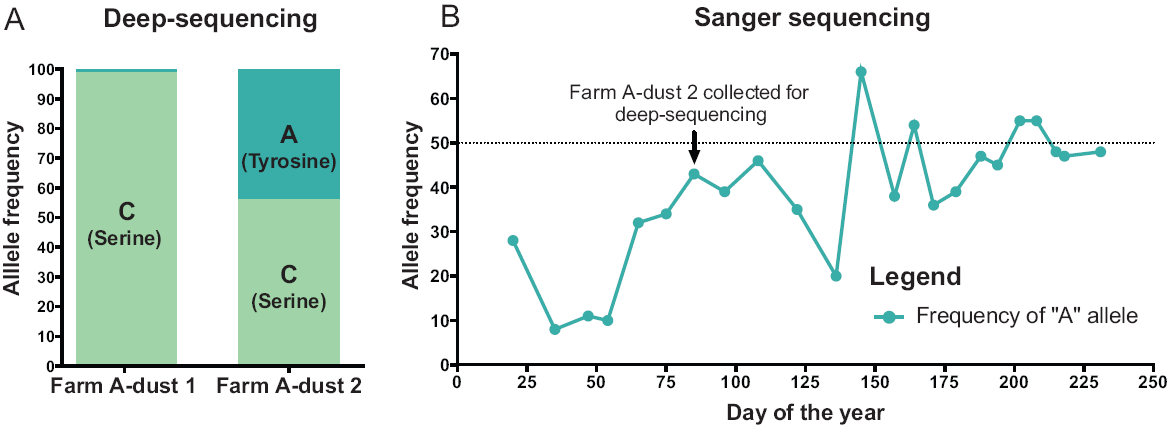
A new polymorphic locus in ICP4, and its shifting allele frequency over time. **(A)** HTS data revealed a new polymorphic locus in ICP4 (MDV084) at nucleotide position 5,495. In the spatially- and temporally-separated dust samples from Farm A (see **Figure 1A** and **Methods** for details), we observed a different prevalence of C (encoding serine) and A (encoding tyrosine) alleles. **(B)** Using targeted Sanger sequencing of this locus, time-separated dust samples spanning nine months were Sanger-sequenced to track polymorphism frequency at this locus over time. The major and minor allele frequencies at this locus varied widely across time, and the major allele switched from C to A more than twice during this time.

Given the very different allele frequencies at this ICP4 locus between two houses on the same farm 11 months apart, we examined dust samples from one of the houses over 9 months with targeted Sanger sequencing of this SNP **(Figure 5B).** We found that this locus was highly polymorphic in time-separated dust samples. The A (Tyrosine) allele rose to almost 50% frequency in the 9 month period. In four of the dust samples, the A (Tyrosine) allele was dominant over the C (serine) allele. This reversible fluctuation in allele frequencies over a short period of time is unprecedented for alphaherpesviruses so far as we know. However, recent studies on human cytomegalovirus (HCMV) have shown that selection can cause viral populations to evolve in short periods of time (34, 35). While this is only one example of a polymorphic locus that shifts in frequency over time, similar approaches could be used at any of the hundreds of polymorphic loci detected here **(Supplemental Table S3).**

### Comparison of field isolates of MDV-1 to previously sequenced isolates

To compare these new field-based MDV genomes to previously sequenced isolates of MDV, we created a multiple sequence alignment of all available MDV-1 genomes (14, 15, 17-21, 47, 48). The multiple sequence alignment was used to generate a dendrogram depicting genetic relatedness (see **Methods).** We observed that the five new isolates form a separate group when compared to all previously sequenced isolates **(Figure 6).** This may result from geographic differences as previously seen for HSV-1 and VZV (27, 49-52), or from temporal differences in the time of sample isolations, or from the lack of cell-culture adaptation in these new genomes.

**Figure 6:**
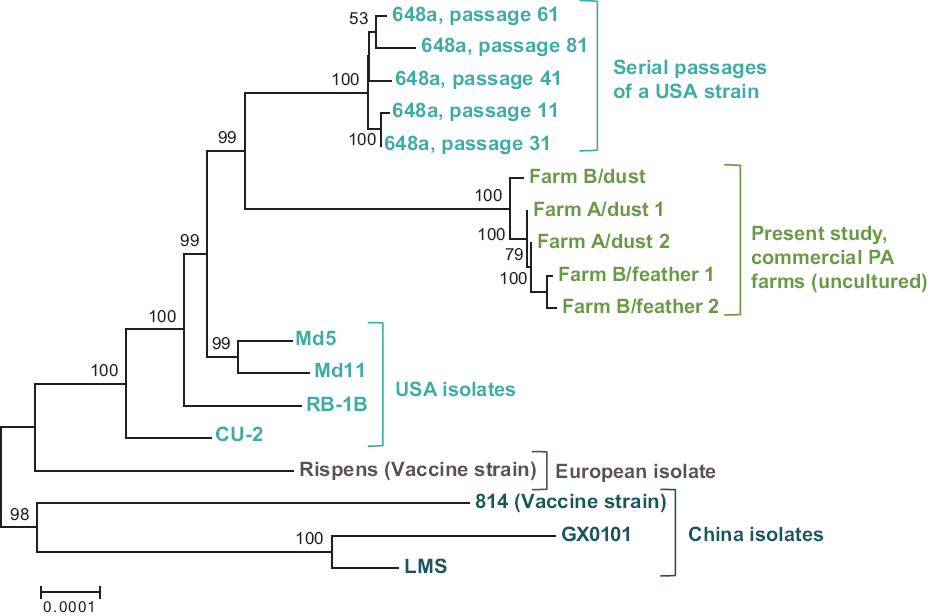
Dendrogram of genetic distances among all sequenced MDV-1 genomes. Using a multiple-genome alignment of all available complete MDV-1 genomes, we calculated the evolutionary distances between genomes using the Jukes-Cantor model. A dendrogram was then created using the neighbor-joining method in MEGA with 1000 bootstraps. The five new field-sampled MDV-1 genomes (green) formed a separate group between the two clusters of USA isolates (blue). The European vaccine strain (Rispens) formed a separate clade, as did the three Chinese MDV-1 genomes (aqua). GenBank Accessions for all strains: new genomes, Table 1; Passage ll-648a, JQ806361; Passage 31-648a, JQ806362; Passage 61-648a, JQ809692; Passage 41-648a, JQ809691; Passage 81-648a, JQ820250; CU-2, EU499381; RB-1B, EF523390; Mdll, 170950; Md5, AF243438; Rispens (CVI988), DQ530348; 814, JF742597; GX0101, JX844666; LMS, JQ314003.

We also noted a distinctive mutation in the genes encoding glycoprotein L (gL; also known as UL1 or MDV013). All of the field isolates had a 12 nucleotide deletion in gL that has been described previously in strains from the Eastern USA. This deletion is found predominantly in very virulent or hypervirulent strains (vv and vv+, in MDV-1 pathotyping nomenclature (1)) (53–56). This deletion falls in the putative cleavage site of gL, which is necessary for its post-translational modification in the endoplasmic reticulum (54). Glycoprotein L forms a complex with another glycoprotein H (gH). The gH/gL dimer is conserved across the *Herpesviridae* family and has been associated with virus entry (57, 58).

These field-isolated genomes also contain a number of previously characterized variations in the oncogenesis-associated Marek’s EcoRI-Q-encoded protein (Meq; also known as MDV005, MDV076, and RLORF7). We observed three substitutions in the C-terminal (transactivation) domain of Meq (P153Q, P176A, P217A) (59). The first two of these variations have been previously associated with MDV-1 strains of very virulent and hypervirulent pathotypes (vv and vv+) (53, 60, 61), while the third mutation has been shown to enhance transactivation (62). In contrast, the field isolates lacked the 59 AA insertion in the Meq proline repeats that is often associated with attenuation, as seen in the vaccine strain CVI988 and the mildly-virulent strain CU-2 (47, 48, 63, 64). We also observed a Cl 19R substitution in all five field-derived genomes, which is absent from attenuated and mildly-virulent isolates. This Cl 19R mutation falls in the LXCXE motif of Meq, which normally binds to the tumor suppressor protein Rb to regulate cell cycle progression (53, 65). Although comprehensive *in vitro* and *in vivo* studies will be required to fully understand the biological implications of these variations, sequence comparisons of both gL and Meq from dust and feather genomes suggest that these closely resemble highly virulent (vv and vv+) variants of MDV-1 (47, 48). This is corroborated by the dendrogram **(Figure 6),** where the dust- and feather-derived genomes cluster closely with 648a, which is a highly virulent (vv+) MDV isolate.

### Assessment of taxonomic diversity in dust and feathers

As noted in **Table 1,** only a fraction of the reads obtained from each sequencing library were specific to MDV-1. We analyzed the remaining sequences to gain insight into the taxonomic diversity found in poultry dust and feathers. Since our enrichment for viral capsids removed most host and environmental contaminants, the taxa observed here represent only a fraction of the material present initially. However it provides useful insight into the overall complexity of each sample type. The results of the classification for Farm B-dust, Farm B-feather 1, and Farm B-feather 2 are shown in **Supplemental Figure S4.** We divided the sequence reads by the different kingdoms they represent. Complete lists of taxonomic diversity for all samples to the family level are listed in **Supplemental Table S5.** As expected, the taxonomic diversity of dust is greater than that of feather samples. The majority of sequences in the dust samples mapped to the chicken genome, and only about 2-5% were MDV specific (see also **Table 1,** line 6). We found that single feathers were a better source of MDV DNA, due to their reduced level of taxonomic diversity and higher percentage of MDV-specific reads **(Table 1,** line 6 and **Supplemental Figure S4).**

## Discussion

This study presents the first description of MDV-1 genomes sequenced directly from a field setting. This work builds on recent efforts to sequence VZV and HCMV genomes directly from human clinical samples, but importantly the approaches presented here do not employ either the oligo-enrichment used for VZV or the PCR-amplicon strategy used for HCMV (10, 33-35, 66). This makes our technique widely accessible and reduces potential methodological bias. It is also more rapid to implement and is applicable to the isolation of unknown large DNA viruses, since it does not rely on sequence-specific enrichment strategies. These five genomes were interrogated at the level of comparing consensus genomes - between-host variation - as well as within each consensus genome - within-host variation. By following up with targeted PCR and Sanger sequencing, we demonstrate that HTS can rapidly empower molecular epidemiological field surveillance of loci undergoing genetic shifts.

Although a limited number of non-synonymous differences were detected between the field samples compared here, it is striking that several of these (vLIP, LORF2, UL42) have been previously demonstrated to have roles in virulence and immune evasion. The N-glycosylated protein viral lipase (vLIP; MDV010) encodes a 120 kDa protein that is required for lytic virus replication in chickens (40, 3). The vLIP gene of MDV-1 is homologous to other viruses in the *Mardivirus* genus as well as to avian adenoviruses (67–69). The S501A mutation in the second exon of vLIP protein is not present in the conserved region that bears homology to other pancreatic lipases (40). The viral protein LORF2 (MDV012) is a viral immune evasion gene that suppresses MHC class I expression by inhibiting TAP transporter delivery of peptides to the endoplasmic reticulum (41). LORF2 is unique to the non-mammalian *Mardivirus* clade, but its function is analogous to that of the mammalian alphaherpesvirus product UL49.5 (41, 70, 71). Another study has shown that LORF2 is an essential phosphoprotein with a potential role as a nuclear/cytoplasmic shuttling protein (72). Interestingly, we also observed a 22 amino acid insertion in the DNA polymerase processivity subunit protein UL42 (MDV055). In herpesviruses, UL42 has been recognized as an integral part of the DNA polymerase complex, interacting directly with DNA and forming a heterodimer with the catalytic subunit of the polymerase (73–75). In HSV-1, the N-terminal two thirds of UL42 have been shown to be sufficient for all known functions of UL42; the insertion in Farm B-dust falls at the edge of this N-terminal region (76). The non-synonymous mutations and insertions detected here warrant further study to evaluate their impacts on protein function and viral fitness *in vivo.* The fact that any coding differences were observed in this small sampling of field-derived genomes suggests that the natural ecology of MDV-1 may include mutations and adaptations in protein function, in addition to genetic drift.

Drug resistance and vaccine failure have been attributed to the variation present in viral populations (10, 33, 77). Polymorphic populations allow viruses to adapt to diverse environments and withstand changing selective pressures, such as evading the host immune system, adapting to different tissue compartments, and facilitating transmission between hosts (10, 26, 33-35, 66, 77, 78). Polymorphisms that were not fully penetrant in the consensus genomes, but that may be fodder for future selection, include residues in genes associated with virulence and immune evasion, such as ICP4 **(Figure 5),** Meq, pp38, vLIP, LORF2, and others **(Supplemental Table S3).** The non-synonymous polymorphism that we observed in Meq is a low-frequency variant present in the C-terminal domain (I201L) (Farm B-dust, **Supplemental Table S3).** However a comparison of 88 different Meq sequences from GenBank and unpublished field isolates (37) did not reveal any examples where leucine was the dominant allele; all sequenced isolates to date have isoleucine at position 201.

Previous studies have examined the accumulation of polymorphic loci in MDV-1 genomes after serial passage *in vitro* (30, 31). Overall, we found a similar quantity of polymorphisms in field-derived genomes as found in these prior studies, but we did not find any specific polymorphic loci that were identical between field-derived and *in* v/Yro-passaged genomes (30, 31). The genes ICP4 (MDV084 / MDV100), LORF2 (MDV012), UL42 and MDV020 contain polymorphic in both field and serially-passaged isolates, albeit at different loci (30, 31). It is noteworthy that these coding variations are detected despite signs of clearance of polymorphisms from coding regions **(Figure 4),** as indicated by the higher-than-expected ratios of intergenic to coding polymorphisms in these genomes. Together these findings suggest that MDV-1 exhibits genetic variation and undergoes rapid selection in the field, which may demonstrate the basis of its ability to overcome vaccine induced host-resistance to infection (3, 79, 80, 5).

For the viral transactivation protein ICP4, we explored the penetrance of a polymorphic locus (nucleotide position 5,495) both in full-length genomes, and also via targeted sequencing over time. Most of the work on this region in the MDV-1 genome has actually focused on the LAT transcripts that lie antisense to the ICP4 gene (44, 45). This polymorphic locus could thus impact either ICP4’s coding sequence (AA 1,832 serine vs. tyrosine) or the sequence of the LAT transcripts. This variation in ICP4 lies in the C-terminal domain, which in HSV-1 has been implicated in the DNA synthesis, late gene expression, and intranuclear localization functions of ICP4 (81, 82). This combination of deep-sequencing genomic approaches to detect new polymorphic loci, and fast gene-specific surveillance to track changes in SNP frequency over a larger number of samples, illustrate the power of high-quality full genome sequences from field samples to provide powerful new markers for field ecology.

Our comparison of new field-isolated MDV-1 genomes revealed a distinct genetic clustering of these genomes, separate from other previously sequenced MDV-1 genomes **(Figure 6).** This pattern may results from geographic and temporal drift in these strains, or from the wild, virulent nature of these strains vs. the adaptation(s) to tissue culture in all prior MDV-1 genome sequences. The impact of geography on the genetic relatedness of herpesvirus genomes has been previously shown for related alphaherpesviruses such as VZV and HSV-1 (27, 49-52). Phenomena such as recombination can also have an impact on the clustering pattern of MDV isolates. It is worth noting that the genetic distance dendrogram constructed here included genomes from isolates that were collected over a 40 year span, which introduces the potential for temporal drift (14, 15, 17-21, 47, 48). Agricultural and farming practices have evolved significantly during this time, and we presume that pathogens have kept pace. To truly understand the global diversity of MDV, future studies will need to include the impacts of recombination and polymorphisms within samples, in addition to the overall consensus-genome differences reflected by static genetic distance analyses.

Prior studies have shown that when MDV is passaged for multiple generations in cell-culture, the virus accumulates a series of mutations, including several that affect virulence (30). The same is true for the betaherpesvirus HCMV (25). Extended passage *in vitro* forms the basis of vaccine attenuation strategies, as for the successful vaccine strain (vOka) of the alphaherpesvirus VZV (83). Cultured viruses can undergo bottlenecks during initial adaptation to cell culture, and they may accumulate variations and loss of function mutations by genetic drift or positive selection. The variations and mutations thus accumulated may have little relationship to virulence and the balance of variation and selection in the field. We thus anticipate that these field-isolated viral genomes more accurately reflect the genomes of wild MDV-1 that are circulating in the field. The ability to access and compare virus from virulent infections in the field will enable future analyses of vaccine-break viruses.

Our data and approaches provide powerful new tools to measure viral diversity in field settings, and to track changes in large DNA virus populations over time in hosts and ecosystems. In the case of MDV-1, targeted surveillance based on an initial genomic survey could be used to track viral spread across a geographic area, or between multiple end-users associated with a single parent corporation **(Figure 1B)**. Similar approaches could be implemented for public-or animal-health programs, for instance to guide management decisions on how to limit pathogen spread and contain airborne pathogens. The ability to sequence and compare large viral genomes directly from individual hosts and field sites will allow a new level of interrogation of host-virus fitness interactions, which form the basis of host resistance to infection **(Figure 1B)**. Finally, the analysis of viral genomes from single feather follicles, as from single VZV vesicles, enables our first insights into naturally-occurring within-host variation during infection and transmission **(Figure 1B).** Evidence from tissue compartmentalization studies in HCMV and VZV suggests that viral genomes differ in distinct body niches (10, 35, 66). These new technique will enable us to ask similar questions about MDV-1, and to begin exploring the relative fitness levels of viruses found in different tissue compartments.

## Materials and Methods

### Collection of dust and feathers

Samples were collected from two commercial-scale farms in central Pennsylvania, where each poultry building housed 25,000-30,000 individuals **(Figure 1A).** The poultry on both farms were the same breed and strain of colored (“red”) commercial broiler chicken from the same hatchery and company. Dust samples were collected into 1.5 ml tubes from fan louvers. This location contains less moisture and contaminants than floor-collected samples, and represents a mixture of air-borne virus particles and feather dander. Sequential samples from Farm A **(Table 1)** were collected 11 months apart, from adjacent houses on the same farm **(Figure 1A)**. Samples from Farm B **(Table 1)** were collected from a single house, at a single point in time. Feathers were collected just before hosts were transported from the farms for sale, to maximize the potential for infection and high viral titer. At the time of collection the animals were 10-12 weeks old. Ten individuals were chosen randomly throughout the entirety of one house for feather collection. Two feathers from each animal were collected from the axillary track (breast feathers). The distal 0.5-1.0 cm proximal shaft or feather tip, which contains the feather pulp, was snipped into a sterile 1.5 ml micro-tube containing a single sterile 5 mm steel bead (Qiagen). On return to the laboratory, tubes were stored at −80°C until processing. One feather from each animal was tested for the presence and quantity and MDV-1 present (see below for quantitative PCR details). The remaining feathers from the two animals with highest apparent MDV-1 titer were used for a more thorough DNA extraction (see below for details) and next-generation sequencing. Animal procedures were approved by the Institutional Animal Care and Use Committee of the Pennsylvania State University (IACUC Protocol#: 46599).

### Viral DNA isolation from dust

MDV nucleocapsids were isolated from dust as indicated in **Supplemental Figure S1A.** Dust collected from poultry houses was stored in 50 ml polypropylene Falcon^®^ tubes (Corning) at 4°C until required. 500 mg of dust was suspended in 6.5 ml of 1X phosphate buffered saline (PBS). To distribute the dust particles into solution and help release cell-associated virus, the mixture was vortexed vigorously until homogenous and centrifuged at 2000 × g for 10 minutes. This supernatant was further agitated on ice for 30 sec. using a Sonica Ultrasonic Processor Q125 (probe sonicator with 1/8^th^ inch microtip) set to 20% amplitude. It was then vortexed before being centrifuged for a further 10 minutes at 2000 × g. To enrich viral capsids away from the remaining contaminants, the supernatant (approximately 5 ml in volume) was subjected to a series of filtration steps. First, we used a Corning^®^ surfactant-free cellulose acetate (SFCA) filter (0.8 µM) that had been soaked overnight in fetal bovine serum (FBS) to remove particles at the level of eukaryotic cells, and bacteria. To remove smaller contaminants, the flow-through was then passed through a Millipore Express^®^ PLUS Membrane vacuum filter (0.22 µM) and the membrane subsequently washed twice with 2.5 ml of PBS. To remove contaminant DNA, the final filtrate (approximately 10 ml in volume) was treated with DNase (Sigma) at a concentration of 0.1 mg/ml for 30 minutes at room temperature. In the absence of DNase treatment we observed a higher yield of viral DNA, but with much lower purity (data not shown). The MDV nucleocapsids present in the DNase-treated solution were captured on a polyethersulfone (PES) membrane (VWR) filter (0.1 µM). This filter membrane trapped the viral nucleocapsids, which are between 0.1-0.2 µm (84). An increased MDV purity, but ultimately reduced total nanograms of DNA yield, may be achieved by washing this membrane once with 2.5 ml PBS (see **Supplemental Table S1).** In the future, samples with a higher percentage of MDV DNA could be obtained by applying these wash steps to all components of the sample pool. The membrane was then carefully excised using a sterile needle and forceps, and laid - exit side downwards - in a sterile 5 cm diameter plastic petri-dish where it was folded twice lengthwise. The “rolled” membrane was then placed into a 2 ml micro-tube containing 1.8 ml of lysis solution (ATL buffer and Proteinase K from the DNeasy^®^ Blood and Tissue kit, Qiagen). Digestion was allowed to proceed at 56°C for 1 hour on an incubating microplate shaker (VWR) set to 1100 rpm. The membrane was then removed, held vertically over a tilted sterile 5 cm diameter plastic petri-dish and washed with a small volume of the lysis solution (from the 2 ml micro-tube). This wash was subsequently returned to the 2 ml micro-tube and the tube replaced on the heated shaker where it was allowed to incubate overnight. The following day, the DNA was isolated as per manufacturer’s instructions using the DNeasy^®^ Blood and Tissue kit (Qiagen). DNA was eluted in 200 µl DNase-free water. Ten to fourteen aliquots of 500 mg each were used to obtain sufficient DNA for each dust sample (see **Supplemental Table S1).** Quantitative PCR was used to assess the copy number of viral genomes in the resulting DNA. Total yield and percent MDV-1 vs. MDV-2 DNA are listed in **Supplemental Table S1.**

### Isolation of viral DNA from feather follicles

The protocol for extraction of MDV DNA from feather follicles was optimized for the smaller input material and an expectation of higher purity **(Supplemental Figure S1B).** Sequential size filters were not used to filter out contaminants from feather follicles, since these direct host samples have fewer impurities than the environmental samples of dust. However, the feather follicle cells were encased inside the keratinaceous shell of the feather tip, which required disruption to release the cells. Each tube containing a single feather tip and one sterile 5 mm diameter steel bead was allowed to thaw, and then 200 µl of PBS was added and the sample bead-beaten for 30 seconds at 30 Hz using a Tissuelyser (Qiagen) **(Supplemental Figure S1B).** Vigorous bead-beating achieved the desired destruction of the follicle tip. To dissociate the cells, 80 µl of 2.5 mg/ml trypsin (Sigma) and 720 µl of PBS were then added (final trypsin concentration: 0.8 mg/ml), and the solution was transferred to a new sterile 2 ml micro-tube and incubated for 2 hours at 37°C on a heated microplate shaker (VWR) set to 700 rpm. To release cell-associated virus, the suspension was then sonicated on ice for 30 seconds using a Sonica Ultrasonic Processor Q125 (probe sonicator with 1/8^th^ inch microtip) set to 50% amplitude. DNase I was added to a final concentration of 0.1 mg/ml and allowed to digest for 1 hour at room temperature to remove non-encapsidated DNA. An equal volume of lysis solution (ATL buffer and Proteinase K from the DNeasy^®^ Blood and Tissue kit, Qiagen) was added and the sample was incubated over night at 56°C on an Incubating Microplate Shaker (VWR) set to 1100 rpm. The following day, the DNA was isolated as per manufacturer’s instructions using the DNeasy^®^ Blood and Tissue kit (Qiagen). While the overall amount of DNA obtained from feather follicles was lower than that obtained from pooled dust samples **(Supplemental Table S2)**, it was of higher purity and was sufficient to generate libraries for sequencing **(Table 1,** lines 1-3).

### Measurement of total DNA and quantification of viral DNA

The total amount of DNA present in the samples was quantified by fluorescence analysis using a Qubit^®^ fluorescence assay (Invitrogen) following the manufacturer’s recommended protocol. MDV genome copy numbers were determined using serotype-specific quantitative PCR (qPCR) primers and probes, targeting either the MDV-1 pp38 (MDV073; previously known as LORF14a) gene or MDV-2 (SB-1 strain) DNA polymerase (UL42, MDV055) gene. The MDV-1 assay was designed by Sue Baigent: forward primer (Spp38for) 5’-GAGCTAACCGGAGAGGGAGA-3’; reverse primer (Spp38rev) 5’-CGCATACCGACTTTCGTCAA-3’; probe (MDV-1) 6FAM-CCCACTGTGACAGCC-BHQ1 (S. Baigent, *pers. comm.).* The MDV-2 assay is that of Islam et al. (85), but with a shorter MGB probe (6FAM-GTAATGCACCCGTGAC-MGB) in place of their BHQ-2 probe. Real-time quantitative PCRs were performed on an ABI Prism 7500 Fast System with an initial denaturation of 95°C for 20 seconds followed by 40 cycles of denaturation at 95°C for 3 seconds and annealing and extension at 60°C for 30 seconds. Both assays included 4 µl of DNA in a total PCR reaction volume of 20 µl with IX PerfeCTa™ qPCR FastMix™ (Quanta Biosciences), forward and reverse primers at 300 nM and TaqMan ^®^ BHQ (MDV-1) or MGB (MDV-2) probes (Sigma and Life Sciences, respectively) at 100 nM and 200 nM, respectively. In addition each qPCR reaction incorporated 2 µl BSA (Sigma). Absolute quantification of genomes was based on a standard curve of serially diluted plasmids cloned from the respective target genes. The absolute quantification obtained was then converted to concentration. Once the concentration of the total DNA, MDV-1, and MDV-2 DNA present in the sample were known, we calculated the percentage of MDV-1 and MDV-2 genomic DNA in the total DNA pool (see **Supplemental Tables S1–S2).**

### Illumina next-generation sequencing

Sequencing libraries for each of the isolates were prepared using the Illumina TruSeq Nano DNA Sample Prep Kit, according to the manufacturer’s recommended protocol for sequencing of genomic DNA. Genomic DNA inputs used for each sample are listed in **Table 1.** The DNA fragment size selected for library construction was 550 base pairs (bp). All the samples were sequenced on an in-house Illumina MiSeq using version 3 sequencing chemistry to obtain paired-end sequences of 300 × 300 bp. Base calling and image analysis was performed with the MiSeq Control Software (MCS) version 2.3.0.

### Consensus genome assembly

As our samples contained DNA from many more organisms than just MDV, we developed a computational workflow **(Supplemental Figure S2)** to preprocess our data prior to assembly. A local BLAST database was created from every *Gallid herpesvirus* genome available in GenBank. All sequence reads for each sample were then compared to this database using BLASTN (86) with a loose e-value less than or equal to 10’^2^ in order to computationally enrich for sequences related to MDV. These “MDV-like” reads were then processed for downstream genome assembly. The use of bivalent vaccine made it possible for us to readily distinguish sequence reads that resulted from the shedding of virulent MDV-1 vs. vaccine virus (MDV-2 or HVT) strains. The overall DNA identity of MDV-1 and MDV-2 is just 61% (87). In a comparison of strains MDV-1 Md5 (NC_002229) and MDV-2 SB-1 (HQ840738), we found no spans of identical DNA greater than 50 bp (data not shown). This allowed us to accurately distinguish these 300 x 300 bp MiSeq sequence reads as being derived from either MDV-1 or MDV-2.

MDV genomes were assembled using the viral genome assembly VirGA (39) workflow which combines quality control preprocessing of reads, *de novo* assembly, genome linearization and annotation, and post-assembly quality assessments. For the reference-guided portion of viral genome assembly in VirGA, the *Gallid herpesvirus 2* (MDV-1) strain MD5 was used (GenBank Accession: NC_002229.3). These new genomes were named according to recent recommendations, as outlined by Kuhn et al (88). We use shortened forms of these names throughout the manuscript (see **Table 1** for short names). The full names for all five genomes are as follows: MDV-1 *Gallus domesticus-*wt/Pennsylvania, USA/2015/Farm A-dust 1; MDV-1 *Gallus domesticus-*wt/Pennsylvania, USA/2015/Farm A-dust 2; MDV-1 *Gallus domesticus-* wt/Pennsylvania, USA/2015/Farm B-dust; MDV-1 *Gallus domesticus-*wt/Pennsylvania, USA/2015/Farm B-feather 1; MDV-1 *Gallus domesticus-*wt/Pennsylvania, USA/2015/Farm B-feather 2. GenBank Accessions are listed below and in **Table 1.** Annotated copies of each genome, in a format compatible with genome- and sequence browsers, are available at the Pennsylvania State University ScholarSphere data repository: https://scholarsphere.psu.edu/collections/1544bpl4i.

### Between-sample: consensus genome comparisons

Clustalw2 (43) was used to construct pairwise global nucleotide alignments between whole genome sequences, and pairwise global amino acid alignments between open reading frames. These alignments were utilized by downstream custom Python scripts to calculate percent identity, protein differences, and variation between samples.

The proline-rich region of UL36 (also known as VP 1/2 or MDV049), which contains an extended array of tandem repeats, was removed from all five consensus genomes prior to comparison. The amount of polymorphism seen in this region of UL36 is driven by fluctuations in the length of these tandem repeats, as has been seen in prior studies with other alphaherpesviruses such as HSV, VZV, and pseudorabies virus (PRV) (32,48-50). Since the length of extended arrays of perfect repeats cannot be precisely determined by *de novo* assembly (22, 23, 26, 27), we excluded this region from pairwise comparisons of genome-wide variation. Genome alignments with and without the UL36 region removed are archived at the ScholarSphere site: https://scholarsphere.psu.edu/collections/1544bpl4i.

### Within-sample: polymorphism detection within each consensus genome

VarScan v2.2.11 (91) was used to detect variants present within each consensus genome. To aid in differentiating true variants from potential sequencing errors (92), two separate variant calling analyses were explored. (10). Our main polymorphism-detection parameters (used in **Figures 3-4** and **Supplemental Tables S3–S4)** were as follows: minimum variant allele frequency ≥0.02; base call quality ≥20; read depth at the position ≥10; independent reads supporting minor allele ≥2. Directional strand bias ≥90% was excluded; a minimum of two reads in opposing directions was required. For comparison and added stringency, we also explored a second set of parameters (used in **Supplemental Figure S3):** minimum variant allele frequency ≥0.05; base call quality ≥20; read depth at the position ≥100; independent reads supporting minor allele ≥5. Directional strand bias ≥80% was excluded. The variants obtained from VarScan were then mapped back to the genome to understand their distribution and mutational impact using SnpEff and SnpSift (93, 94). Polymorphisms in the proline-rich region of UL36 were excluded, as noted above.

### Testing for signs of selection acting on polymorphic viral populations

For each of our five consensus genomes, which each represent a viral population, we classified the polymorphisms detected into categories of synonymous, non-synonymous, genic-untranslated, or intergenic, based on where each polymorphism was positioned in the genome. For these analyses **(Figure 4),** we were only able to include polymorphisms detected in the three dust genomes, since the total number of polymorphisms obtained from feather genomes was too low for chi-square analysis. First, we calculated the total possible number of single nucleotide mutations that could be categorized as synonymous, non-synonymous, genic-untranslated or intergenic. To remove ambiguity when mutations in overlapping genes could be classified as either synonymous or non-synonymous, genes with alternative splice variants or overlapping reading frames were excluded from these analyses. This removed 25 open reading frames (approximately 21% of the genome). These tallies of potential mutational events were used to calculate the expected fraction of mutations in each category. We preformed chi-squared tests on each dataset to assess whether the observed distribution of polymorphisms matched the expected distribution. We also performed a similar analysis in pairwise fashion **(Supplemental Table S4),** to assess whether the fraction of variants differed from what would be expected by random chance. Pairwise combinations included the following: synonymous vs. non-synonymous, synonymous vs. intergenic, synonymous vs. genic-untranslated, non-synonymous vs. intergenic, non-synonymous vs. genic-untranslated, and intergenic vs. genic-untranslated. Statistically significant outcomes would suggest that recent or historical selection differed between those categories of variants.

### Sanger sequencing of polymorphic locus in ICP4

A potential locus of active selection within the ICP4 (MDV084 / MDV100) gene was detected during deep-sequencing of Farm B-dust. This locus was examined using Sanger sequencing. An approximately 400 bp region of the ICP4 gene was amplified using a Taq PCR Core Kit (Qiagen) and the following primers at 200 nM: forward primer (ICP4selF) 5’AACACCTCTTGCCATGGTTC 3’; reverse primer (ICP4selR) 5’GGACCAATCATCCTCTCTGG 3’. Cycling conditions included an initial denaturation of 95°C for 2 minutes, followed by 40 cycles of denaturation at 95°C for 30 seconds, annealing at 55°C for 30 seconds and extension at 72°C for 1 minute, with a terminal extension at 72°C for 10 minutes. The total reaction volume of 50 µl included 10 µl of DNA and 4 µl BSA (final concentration 0.8 mg/ml). Amplification products were visualized on a 1.5% agarose gel, the target amplicon excised and then purified using the E.Z.N. A. Gel Extraction Kit (Omega Bio-tek). Sanger sequencing was performed by the Penn State Genomics Core Facility utilizing the same primers as used for DNA amplification. The relative peak height of each base call at the polymorphic position was analyzed using the ab1PeakReporter tool (95).

### Genetic distance and dendrogram

Multiple sequence alignments of complete MDV-1 *(Gallid herpesvirus 2)* genomes from GenBank and those assembled by our lab were generated using MAFFT (96). The evolutionary distances were computed using the Jukes-Cantor method (97) and the evolutionary history was inferred using the neighbor-joining method (98) in MEGA6 (99), with 1,000 bootstrap replicates (100). Positions containing gaps and missing data were excluded. The 18-strain genome alignment is archived at ScholarSphere: https://scholarsphere.psu.edu/collections/1544bp14j.

### Taxonomic estimation of non-MDV sequences in dust and feathers

All sequence reads from each sample were submitted to a quality control preprocessing method to remove sequencing primers, artifacts, and areas of low confidence (39). Sequence annotation was performed using a massively iterative all-vs.-all BLASTN (E-value ≤ 10^-2^) approach using the all-nucleotide-database from NCBI. Only a portion of the total sequence read pool could be identified with confidence using this method. We then used *de novo* assembly to extend the length of these unidentified sequences, therefore elongating them into contigs. These were iterated through BLASTN again, which revealed alignment to repetitive regions of the *Gallus domesticus* (chicken) genome. Since the viral DNA enrichment procedures include a level of stochasticity in removal of host and environmental contaminants, the proportion of taxa present is not a definitive outline of those present initially. The results of these classifications are shown in **Supplemental Figure S4** and listed in **Supplemental Table S5.**

### GenBank accession numbers and availability of materials

GenBank Accessions are listed here and in **Table 1:** Farm A - dust 1, KU173116; Farm A - dust 2, KU173115; Farm B - dust, KU173119; Farm B - feather 1, KU173117; Farm B - feather 2, KU173118. Additional files used in this manuscript, such as multiple-sequence alignments of these genomes, are archived and available at ScholarSphere: https://scholarsphere.psu.edu/collections/1544bpl4i

## Acknowledgements

We thank Sue Baigent, Michael DeGiorgio, Peter Kerr and members of Szpara and Read labs for helpful feedback and discussion. This work was supported and inspired by the Center for Infectious Disease Dynamics and the Huck Institutes for the Life Sciences, as well as by startup funds (MLS) from the Pennsylvania State University. This work was part funded by the Institute of General Medical Sciences, National Institutes of Health (R01GM105244; AFR) as part of the joint NSF-NIH-USDA Ecology and Evolution of Infectious Diseases program.

## Supplemental Material Legends and Descriptions

**Supplemental Figure S1. Procedures for enrichment and isolation of MDV DNA from dust or individual feather follicles. (A)** Procedure for enrichment of MDV DNA using dust as the source of viral DNA. Vortexing, centrifugation and sonication were essential to release cell-associated virus into the solution. The virus-containing supernatant was then passed through 0.8 µM and 0.22 µM filters for removal of larger contaminants. The flow-thorough was treated with DNase and the viral particles were captured using 0.1 µM filter. The membrane of the 0.1 µM filter was then excised and used for extraction of the viral DNA. **(B)** Procedure for enrichment of MDV DNA using chicken feather follicle as the source of viral DNA. A feather was mechanically disrupted (bead-beating) and treated with trypsin to break open host cells and release cell-free virus into the solution. The sample was then treated with DNase to remove contaminant DNA. Finally, the viral capsids were lysed to obtain viral genomic DNA.

**Supplemental Figure S2. Workflow for computational enrichment for MDV sequences and subsequent viral genome assembly and taxonomic profiling.** The VirGA workflow (39) requires an input of high-quality HTS data from the viral genome of interest. For this study we added an additional step that selected MDV-like sequence reads from the milieu of dust and feather samples. The sequence reads of interest were obtained by using BLAST to compare all reads against a custom MDV database with an E-value of 10^-2^; these were then submitted to VirGA for assembly. Taxonomic profiling followed a similar path using NCBI’s all-nucleotide database to identify the taxonomic kingdom for each sequence read. In this workflow diagram, parallelograms represent data outputs while rectangles represent computational actions.

**Supplemental Figure S3. Genome-wide distribution of polymorphisms within each consensus genome, using high-stringency criteria.** Polymorphic base calls from each MDV genome were grouped by position in bins of 5 kb and the sum of polymorphisms in each bin was plotted. Stricter parameters of polymorphism detection (see **Methods** for details) revealed a similar distribution to those in **Figure 3.** No polymorphisms were detected in feather-derived genomes using high-stringency criteria, due to their lower coverage depth (see **Table** 1). Note that the upper and lower segments of the y-axis have different scales; the number of polymorphic bases per segment for the split column on the right are thus labeled on the graph.

**Supplemental Figure S4. Taxonomic diversity in dust and chicken feathers from Farm B.** We used an iterative BLASTN workflow to generate taxonomic profiles for all samples from Farm B (see **Methods** for details). Major categories are shown here, with a full list of taxa (to family level) in **Supplemental Table S5.** Farm B-feather 1 and Farm B-feather 2 show less overall diversity, as would be expected from direct host-sampling, vs. the environmental mixture of the dust samples. Since the viral DNA enrichment procedures remove variable amounts of host and environmental contaminants, the proportion of taxa present is representative but not fully descriptive of those present initially. The asterisk indicates sequences that were unclassified or at low prevalence.

**Supplemental Table S1.** [PDF]: Yield and percent MDV 1+MDV2 and total nanograms of DNA in each sample for Farm A-dust 1, Farm A-dust 2, and Farm B-dust

| Samples | Washes on<br>0.1 µm<br>filter<sup>a</sup> | %MDV-1 | %MDV-2 | % MDV1 +<br>MDV2 | DNA (ng) |
| --- | --- | --- | --- | --- | --- |
| **Farm A-dust1** |  |  |  |  |  |
| 1 | 0 | 2.88 | 5.44 | 8.3 | 6.94 |
| 2 | 0 | 2.03 | 5.12 | 7.2 | 6.59 |
| 3 | 0 | 4.16 | 8.39 | 12.5 | 6.73 |
| 4 | 0 | 2.51 | 4.73 | 7.2 | 4.71 |
| 5 | 0 | 1.66 | 3.3 | 4.96 | 6.97 |
| **6** | **1** | **9.13** | **13.99** | **23.12** | **2.69** |
| **7** | **1** | **9.29** | **15.7** | **24.99** | **2.16** |
| **8** | **1** | **5.86** | **10.91** | **16.77** | **3.36** |
| 9 | 0 | 1.89 | 2.98 | 4.9 | 9.81 |
| 10 | 0 | 1.76 | 2.9 | 4.7 | 17.35 |
| 11 | 0 | 2.69 | 5.33 | 8.02 | 8.96 |
| 12 | 0 | 4.49 | 7.8 | 12.29 | 4.14 |
| 13 | 0 | 1.16 | 2.49 | 3.65 | 20 |
| 14 | 0 | 1.36 | 2.83 | 4.19 | 19.47 |
| **Farm A-dust2** |  |  |  |  |  |
| 1 | 0 | 1.5 | 3.16 | 4.66 | 10.69 |
| 2 | 0 | 2.55 | 5.62 | 8.17 | 7.18 |
| 3 | 0 | 1.36 | 3.68 | 5.04 | 7.62 |
| 4 | 0 | 1.38 | 2.94 | 4.32 | 9.84 |
| **5** | **1** | **2.71** | **6.19** | **8.9** | **4.11** |
| **6** | **1** | **3.08** | **5.87** | **8.95** | **4.37** |
| **7** | **1** | **2.68** | **4.91** | **7.59** | **5.88** |
| **8** | **1** | **3.49** | **6.24** | **9.73** | **4.88** |
| **9** | **1** | **4.09** | **7.94** | **12.03** | **2.66** |
| **10** | **1** | **6.42** | **10.52** | **16.94** | **3.15** |
| 11 | 0 | 0.26 | 0.91 | 1.17 | 20.35 |
| 12 | 0 | 0.19 | 0.56 | 0.75 | 26.09 |
| 13 | 0 | 0.24 | 0.93 | 1.17 | 15.13 |
| 14 | 0 | 0.36 | 1.21 | 1.57 | 5.62 |
| **Farm B-dust** |  |  |  |  |  |
| 1 | 0 | 0.84 | 6.68 | 7.52 | 14.1 |
| 2 | 0 | 0.46 | 5.2 | 5.66 | 26.64 |
| 3 | 0 | 0.65 | 4.85 | 5.5 | 19.43 |
| 4 | 0 | 0.75 | 5.91 | 6.66 | 16.84 |
| 5 | 0 | 0.23 | 3.67 | 3.9 | 25.9 |
| 6 | 0 | 0.53 | 4.65 | 5.18 | 23.5 |
| **7** | **1** | **1.1** | **14.5** | **15.6** | **4.59** |
| **8** | **1** | **0.95** | **15.77** | **16.72** | **4.29** |
| **9** | **1** | **0.95** | **14.4** | **15.35** | **4.81** |
| **10** | **1** | **1.02** | **10.69** | **11.71** | **3.59** |

<sup>a</sup>Samples that were washed before lysis (bold) yielded a higher percent MDV DNA, but less overall DNA

**Supplemental Table S2.** [PDF]: Yield and percent MDV1+MDV2 and total nanograms of DNA in each sample for Farm B-feathers

| Samples | % MDV-1 | % MDV-2 | % MDV-1<br>+MDV-2 | DNA (ng) |
| --- | --- | --- | --- | --- |
| Feather 1 | 40.59 | 0.12 | 40.72 | 11.97 |
| Feather 2 | 5.68 | 0.02 | 5.70 | 27.36 |

**Supplemental Table S3.**
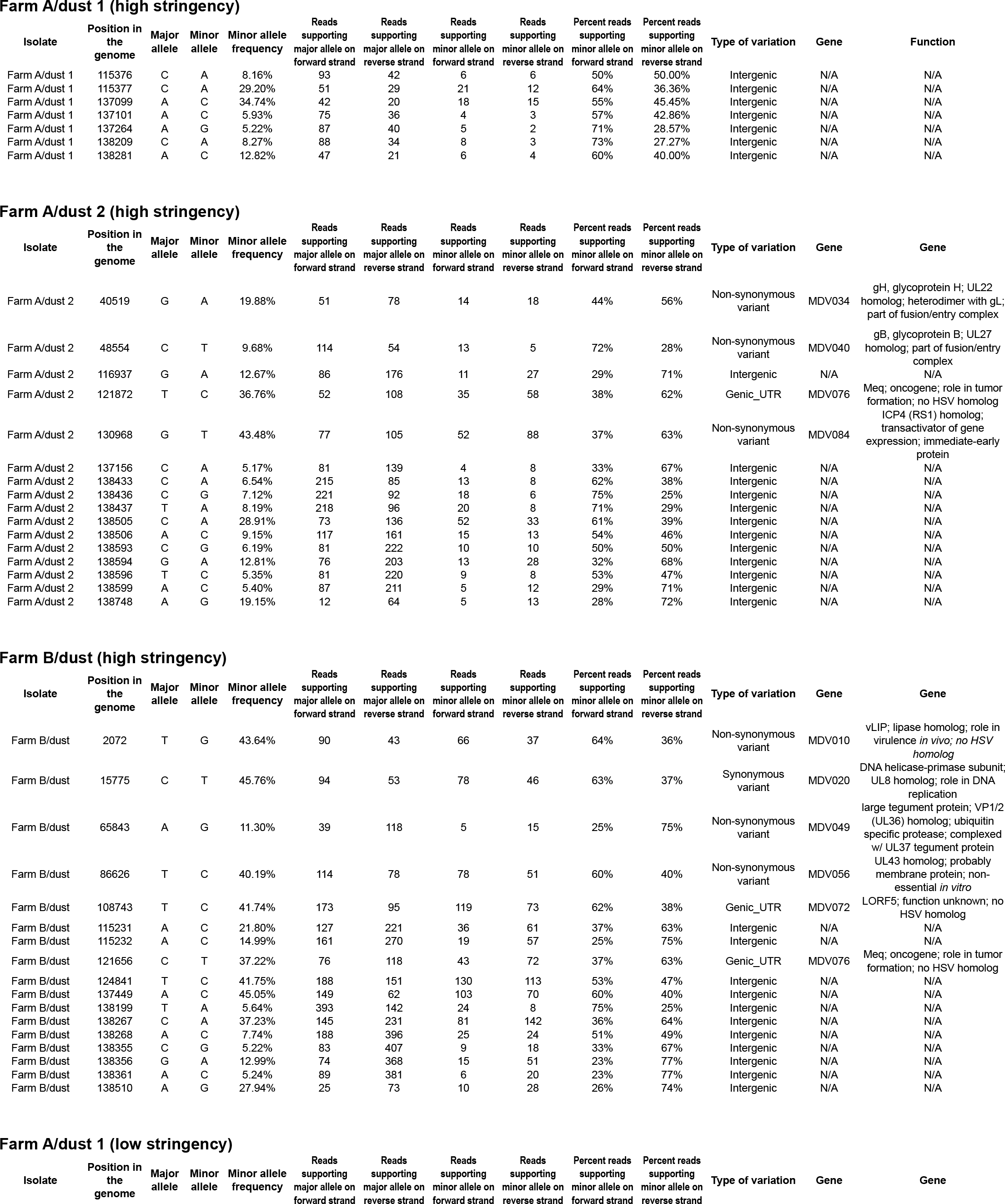
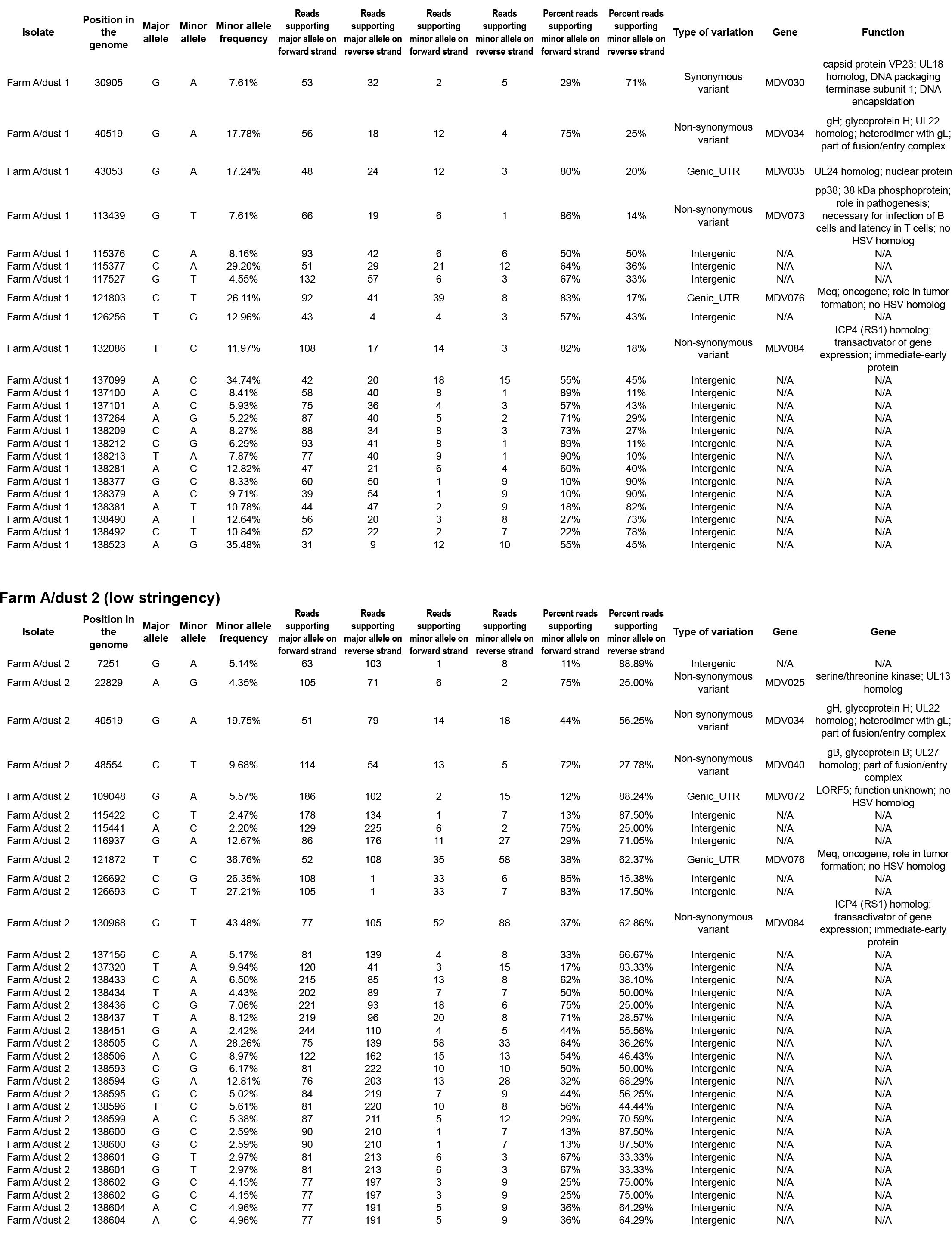
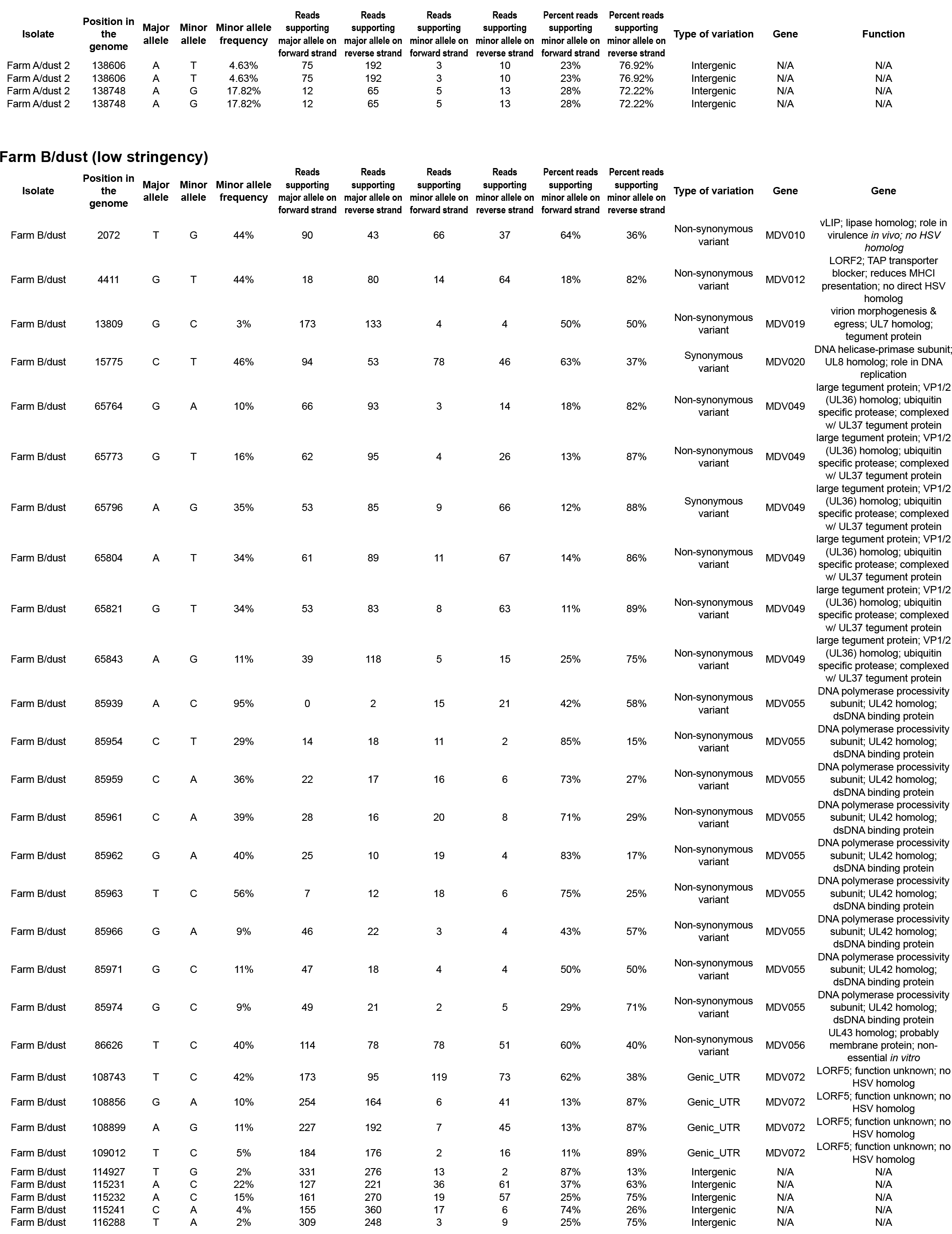
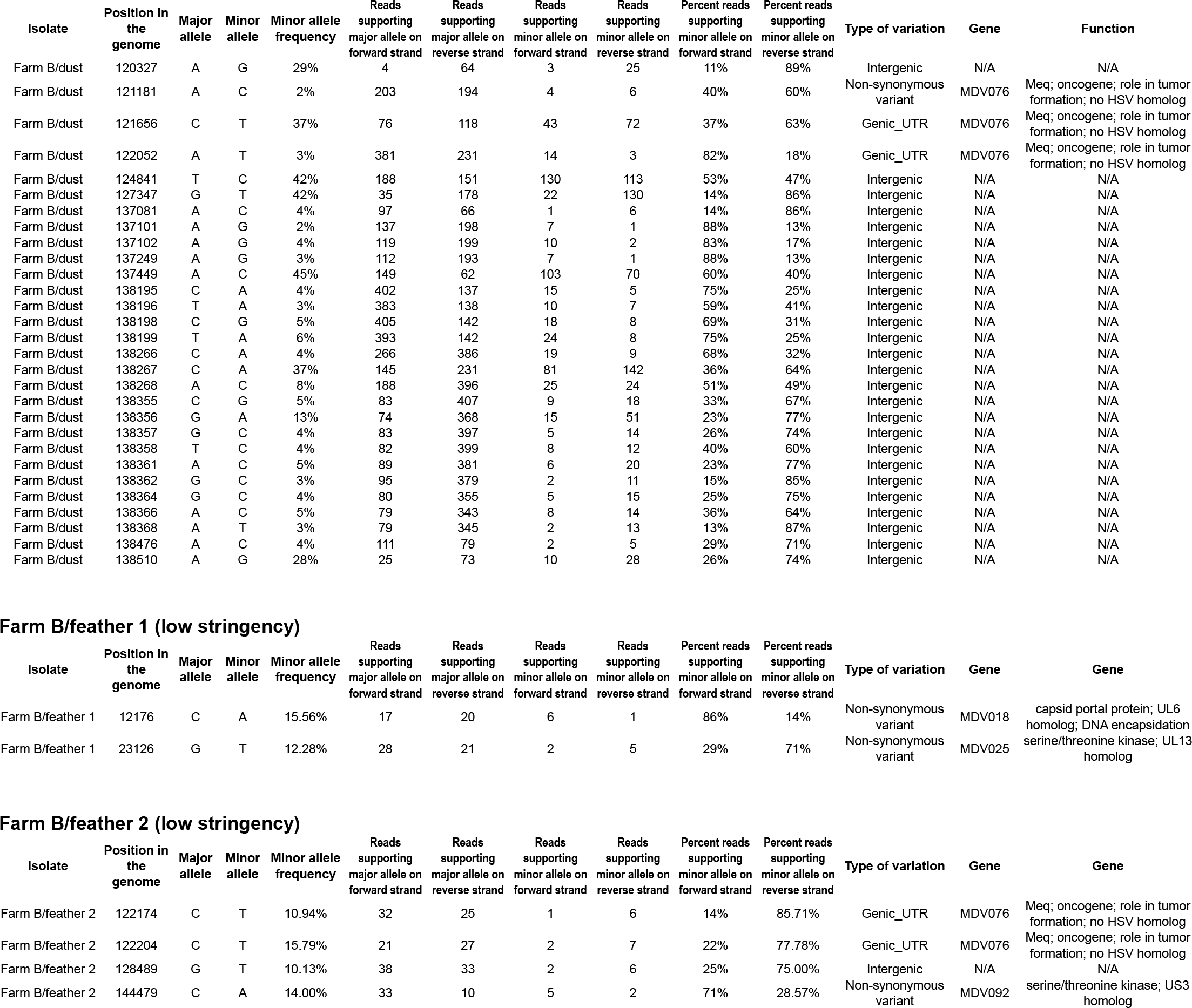
[Excel]: Summary and annotation of all polymorphic loci detected in MDV-1 consensus genomes.

**Supplemental Table S4.** [PDF]: Chi-squared values from pairwise comparisons of different categories of polymorphisms.

| **Sample<sup>a</sup>** | **Intergenic vs. synonymous** | **Intergenic vs. non-synonymous** | **Intergenic vs. genic untranslated** | **Synonymous vs. non-synonymous** | **Synonymous vs. genic untranslated** | **Non-synonymous vs. genic untranslated** |
| --- | --- | --- | --- | --- | --- | --- |
| **Farm A-dust 1** | $\chi^2=16.6$<br>(p = <0.001) | $\chi^2=55.47$<br>(p = <0.001) | $\chi^2=3.74$<br>(p = 0.053) | $\chi^2=0.03$<br>(p=0.873) | $\chi^2=0.83$<br>(p = 0.361) | $\chi^2=1.73$<br>(p = 0.189) |
| **Farm A-dust 2** | $\chi^2=31.76$<br>(p = <0.001) | $\chi^2=94.93$<br>(p = <0.001) | $\chi^2=9.48$<br>(p = 0.002) | $\chi^2=1.11$<br>(p = 0.292) | $\chi^2=2.72$<br>(p = 0.099) | $\chi^2=0.69$<br>(p = 0.407) |
| **Farm B-dust** | $\chi^2=25.27$<br>(p = <0.001) | $\chi^2=47.32$<br>(p = <0.001) | $\chi^2=5.39$<br>(p = 0.020) | $\chi^2=1.83$<br>(p = 0.176) | $\chi^2=1.61$<br>(p = 0.205) | $\chi^2=0.09$<br>(p = 0.759) |

<sup>a</sup>Degrees of freedom (d.f.) = 1 for all comparisons; p indicates p-value.

**Supplemental Table S5.** [Excel]: Summary of taxonomic classification to the family level for all samples

| Family Name | Farm B/dust | Farm B/feather 1 | Farm B/feather 2 | Farm A/dust 1 | Farm A/dust 2 |
| --- | --- | --- | --- | --- | --- |
| Herpesviridae | 1,372,838 | 103,939 | 165,216 | 370,393 | 512,531 |
| Chicken | 19,119,306 | 212,314 | 121,060 | 11,783,342 | 21,215,016 |
| <100 | 570,555 | 50,009 | 27,895 | 794,467 | 533,986 |
| Gramineae | 48,512 | 6,145 | 13,189 | 17,909 | 59,709 |
| Meleagrididae | 47,292 | 9,570 | 6,352 | 16,493 | 76,814 |
| Methylobacteriaceae | 35,567 | 2,660 | 5,205 | 356 | 58,844 |
| other sequences | 27,430 | 1,678 | 3,012 | 10,400 | 14,568 |
| Propionibacteriaceae | 15,405 | 0 | 148 | 6,197 | 12,459 |
| Bovidae | 1,131,177 | 0 | 0 | 8,055 | 39,873 |
| Bradyrhizobiaceae | 986,082 | 0 | 0 | 682 | 99,710 |
| Dermabacteraceae | 605,631 | 0 | 0 | 75,916 | 375,290 |
| Staphylococcaceae | 337,057 | 0 | 0 | 178,747 | 240,507 |
| Corynebacteriaceae | 229,208 | 0 | 0 | 52,055 | 129,345 |
| unclassified | 196,911 | 0 | 0 | 147,502 | 109,899 |
| Lactobacillaceae | 196,157 | 0 | 0 | 147,351 | 195,604 |
| Babesiidae | 141,039 | 0 | 0 | 3,930 | 10,966 |
| Bacillaceae | 104,791 | 0 | 0 | 77,052 | 97,930 |
| Vira | 102,266 | 0 | 0 | 7,700 | 31,685 |
| Sphingobacteriaceae | 77,220 | 0 | 0 | 36,775 | 68,077 |
| Bacteroidaceae | 72,040 | 0 | 0 | 32,391 | 64,454 |
| Streptococcaceae | 63,088 | 0 | 0 | 23,633 | 42,208 |
| Actinoplanaceae | 55,273 | 0 | 0 | 19,379 | 52,946 |
| Lachnospiraceae | 54,841 | 0 | 0 | 35,767 | 100,136 |
| Siphoviridae | 54,286 | 0 | 0 | 28,164 | 74,766 |
| Clostridiaceae | 53,824 | 0 | 0 | 18,827 | 43,915 |
| biota | 53,379 | 0 | 0 | 20,321 | 64,013 |
| Peptostreptococcaceae | 49,713 | 0 | 0 | 4,585 | 29,781 |
| Ruminococcaceae | 36,850 | 0 | 0 | 22,980 | 63,335 |
| Enterobacteraceae | 35,496 | 0 | 0 | 11,304 | 72,340 |
| Micrococcaceae | 30,584 | 0 | 0 | 13,502 | 28,318 |
| Nocardiaceae | 26,664 | 0 | 0 | 6,869 | 20,515 |
| Myoviridae | 25,744 | 0 | 0 | 26,447 | 56,563 |
| Actinosynnemataceae | 24,644 | 0 | 0 | 7,354 | 21,047 |
| Enterococcaceae | 24,224 | 0 | 0 | 13,055 | 23,442 |
| Mycobacteriaceae | 23,366 | 0 | 0 | 6,151 | 18,818 |
| Pseudomonadaceae | 21,552 | 0 | 0 | 2,478 | 11,341 |
| Burkholderiaceae | 21,347 | 0 | 0 | 0 | 5,960 |
| Rhizobiaceae | 16,069 | 0 | 0 | 103 | 4,745 |
| Nocardiopsaceae | 14,995 | 0 | 0 | 11,804 | 22,330 |
| Sphingomonadaceae | 14,945 | 0 | 0 | 407 | 15,444 |
| Campylobacter group | 13,811 | 0 | 0 | 6,182 | 5,466 |
| Porphyromonadaceae | 13,008 | 0 | 0 | 6,759 | 25,153 |
| Comamonadaceae | 11,483 | 0 | 0 | 623 | 6,288 |
| Rikenellaceae | 11,272 | 0 | 0 | 13,944 | 61,763 |
| Cellulomonadaceae | 10,982 | 0 | 0 | 3,894 | 9,435 |
| Xanthobacteraceae | 10,979 | 0 | 0 | 389 | 2,367 |
| Rhodospirillaceae | 10,498 | 0 | 0 | 499 | 3,578 |
| Microbacteriaceae | 9,978 | 0 | 0 | 3,217 | 8,655 |
| Cervidae | 9,471 | 0 | 0 | 0 | 0 |
| Lysobacteraceae | 9,311 | 0 | 0 | 1,026 | 5,336 |
| Phyllobacteriaceae | 9,070 | 0 | 0 | 217 | 2,283 |
| Promicromonosporaceae | 8,756 | 0 | 0 | 2,975 | 7,802 |
| Dermacoccaceae | 8,713 | 0 | 0 | 2,667 | 7,232 |
| Bifidobacteriaceae | 8,667 | 0 | 0 | 5,241 | 52,059 |
| Coriobacteriaceae | 7,818 | 0 | 0 | 6,884 | 12,377 |
| Geodermatophilaceae | 7,635 | 0 | 0 | 2,153 | 6,594 |
| Paenibacillaceae | 6,921 | 0 | 0 | 4,357 | 7,627 |
| Rhodobacteraceae | 6,764 | 0 | 0 | 331 | 3,802 |
| Caulobacteraceae | 6,531 | 0 | 0 | 685 | 2,499 |
| Listeriaceae | 6,379 | 0 | 0 | 6,555 | 8,296 |
| Nocardioidaceae | 6,376 | 0 | 0 | 2,592 | 5,286 |
| Gordoniaceae | 6,318 | 0 | 0 | 1,779 | 4,944 |
| Pasteurellaceae | 6,071 | 0 | 0 | 2,431 | 1,853 |
| Eubacteriaceae | 5,647 | 0 | 0 | 3,392 | 8,423 |
| Flavobacteriaceae | 5,492 | 0 | 0 | 3,837 | 4,432 |
| Erysipelothrix group | 5,484 | 0 | 0 | 3,451 | 8,925 |
| Alcaligenaceae | 5,223 | 0 | 0 | 561 | 2,550 |
| Frankiaceae | 5,149 | 0 | 0 | 2,071 | 4,817 |

**Supplemental table S5: Summary of classification to the family level for all samples**
|  |  |  |  |  |  |
| --- | --- | --- | --- | --- | --- |
| Peptococcaceae | 4,819 | 0 | 0 | 3,381 | 6,319 |
| Beutenbergiaceae | 4,723 | 0 | 0 | 1,786 | 4,091 |
| Sanguibacteraceae | 4,681 | 0 | 0 | 1,619 | 3,950 |
| Carnobacteriaceae | 4,362 | 0 | 0 | 3,804 | 4,445 |
| Kineosporiaceae | 4,278 | 0 | 0 | 1,351 | 4,031 |
| Catenulisporaceae | 4,007 | 0 | 0 | 973 | 2,611 |
| Megasphaera group | 3,758 | 0 | 0 | 2,066 | 4,526 |
| Rhodocyclaceae | 3,754 | 0 | 0 | 484 | 2,155 |
| Caryophanaceae | 3,378 | 0 | 0 | 2,287 | 3,492 |
| Oscillospiraceae | 3,268 | 0 | 0 | 2,227 | 5,981 |
| Delphinidae | 3,118 | 0 | 0 | 0 | 0 |
| Intrasporangiaceae | 3,105 | 0 | 0 | 822 | 2,230 |
| Verrucomicrobia subdivision 1 | 2,960 | 0 | 0 | 134 | 2,274 |
| Aspergillaceae | 2,918 | 0 | 0 | 0 | 0 |
| Thermoanaerobacterales Family III. Ir | 2,695 | 0 | 0 | 1,907 | 2,999 |
| Acetobacteraceae | 2,675 | 0 | 0 | 0 | 1,164 |
| Leuconostoc group | 2,675 | 0 | 0 | 2,382 | 3,363 |
| Oxalobacteraceae | 2,523 | 0 | 0 | 154 | 1,090 |
| Ancylobacter group | 2,446 | 0 | 0 | 0 | 977 |
| Microsphaeraceae | 2,427 | 0 | 0 | 773 | 2,196 |
| Desulfovibrionaceae | 2,370 | 0 | 0 | 605 | 3,042 |
| Actinomycetaceae | 2,328 | 0 | 0 | 1,052 | 1,947 |
| Tsukamurellaceae | 2,252 | 0 | 0 | 585 | 1,746 |
| Fusobacteriaceae | 2,232 | 0 | 0 | 872 | 1,807 |
| Acinetobacteraceae | 2,138 | 0 | 0 | 10,446 | 4,512 |
| Borrelomycetaceae | 2,006 | 0 | 0 | 546 | 1,011 |
| Cytophaga-Flexibacter group | 1,999 | 0 | 0 | 490 | 1,636 |
| Alcanivorax/Fundibacter group | 1,945 | 0 | 0 | 484 | 1,729 |
| Sapromycetaceae | 1,943 | 0 | 0 | 1,103 | 1,606 |
| Spirochaetaceae | 1,866 | 0 | 0 | 1,069 | 3,796 |
| Muridae | 1,847 | 0 | 0 | 1,525 | 3,953 |
| Ectothiorhodospira group | 1,784 | 0 | 0 | 121 | 1,441 |
| Deinococcaceae | 1,768 | 0 | 0 | 733 | 1,637 |
| Nymphalidae | 1,764 | 0 | 0 | 0 | 112 |
| Glycomycetaceae | 1,732 | 0 | 0 | 575 | 1,515 |
| Thermoanaerobacteraceae | 1,659 | 0 | 0 | 850 | 2,119 |
| Prevotellaceae | 1,573 | 0 | 0 | 244 | 2,229 |
| Aeromonadaceae | 1,534 | 0 | 0 | 273 | 1,471 |
| Anaeromyxobacteraceae | 1,496 | 0 | 0 | 326 | 1,337 |
| Myxococcaceae | 1,458 | 0 | 0 | 443 | 1,417 |
| Microviridae | 1,430 | 0 | 0 | 0 | 758 |
| Geobacteraceae | 1,371 | 0 | 0 | 383 | 1,653 |
| Peptoniphilaceae | 1,357 | 0 | 0 | 820 | 1,317 |
| Chromobacteriaceae | 1,305 | 0 | 0 | 398 | 943 |
| Leptotrichiaceae | 1,302 | 0 | 0 | 674 | 1,496 |
| Sorangiaceae | 1,280 | 0 | 0 | 145 | 943 |
| Chromatiaceae | 1,264 | 0 | 0 | 127 | 963 |
| Rhodobiaceae | 1,239 | 0 | 0 | 0 | 245 |
| Spiroplasmataceae | 1,234 | 0 | 0 | 824 | 1,207 |
| Beijerinckiaceae | 1,198 | 0 | 0 | 0 | 338 |
| Halanaerobiaceae | 1,153 | 0 | 0 | 605 | 936 |
| Suidae | 1,106 | 0 | 0 | 294 | 1,088 |
| Thermaceae | 1,089 | 0 | 0 | 220 | 1,568 |
| Haloarchaeaceae | 1,084 | 0 | 0 | 0 | 215 |
| Sporolactobacillaceae | 1,084 | 0 | 0 | 885 | 1,221 |
| Conexibacteraceae | 1,070 | 0 | 0 | 235 | 832 |
| Brachyspiraceae | 1,065 | 0 | 0 | 572 | 1,003 |
| Acidobacteriaceae | 1,001 | 0 | 0 | 0 | 887 |
| Acidimicrobiaceae | 980 | 0 | 0 | 236 | 478 |
| Cercopithecidae | 882 | 0 | 0 | 600 | 1,146 |
| Hominidae | 849 | 0 | 0 | 39,199 | 3,817 |
| Nolanaceae | 847 | 0 | 0 | 4,110 | 926 |
| Acidaminococcaceae | 818 | 0 | 0 | 515 | 1,206 |
| Thermotogaceae | 794 | 0 | 0 | 456 | 624 |
| Chlorobiaceae | 774 | 0 | 0 | 0 | 954 |
| Clostridiales Family XVIII. Incertae Se | 771 | 0 | 0 | 375 | 1,055 |
| Dietziaceae | 755 | 0 | 0 | 129 | 338 |
| Methanobacteriaceae | 750 | 0 | 0 | 180 | 114 |
| Methylocystaceae | 739 | 0 | 0 | 0 | 202 |
| Aerococcaceae | 716 | 0 | 0 | 531 | 631 |

**Supplemental table S5: Summary of classification to the family level for all samples**
|  |  |  |  |  |  |
| --- | --- | --- | --- | --- | --- |
| Jonesiaceae | 675 | 0 | 0 | 377 | 574 |
| Helicobacteraceae | 671 | 0 | 0 | 0 | 393 |
| Clostridiales Family XVII. Incertae Se | 664 | 0 | 0 | 313 | 949 |
| Cyprinidae | 650 | 0 | 0 | 108 | 174 |
| Methanomassiliicoccaceae | 632 | 0 | 0 | 435 | 690 |
| Haliangiaceae | 628 | 0 | 0 | 146 | 435 |
| Clostridiales Family XV. Incertae Sedi | 626 | 0 | 0 | 244 | 812 |
| Eremotheciaceae | 622 | 0 | 0 | 146 | 312 |
| Brucellaceae | 614 | 0 | 0 | 105 | 185 |
| Podoviridae | 570 | 0 | 0 | 262 | 2,117 |
| Giraffidae | 569 | 0 | 0 | 0 | 0 |
| Alteromonadaceae | 568 | 0 | 0 | 113 | 469 |
| Mustelidae | 568 | 0 | 0 | 0 | 0 |
| Rubrobacteraceae | 548 | 0 | 0 | 118 | 556 |
| Leporidae | 506 | 0 | 0 | 103 | 129 |
| Mimiviridae | 500 | 0 | 0 | 0 | 309 |
| Alicyclobacillaceae | 498 | 0 | 0 | 122 | 564 |
| Equidae | 494 | 0 | 0 | 0 | 0 |
| Opitutaceae | 492 | 0 | 0 | 109 | 524 |
| Euphorbiaceae | 483 | 0 | 0 | 0 | 0 |
| Hyphomonadaceae | 472 | 0 | 0 | 0 | 153 |
| Alcanivoracaceae | 471 | 0 | 0 | 132 | 289 |
| Halobacteroidaceae | 465 | 0 | 0 | 408 | 648 |
| Archangiaceae | 458 | 0 | 0 | 149 | 372 |
| Sphaerobacteraceae | 458 | 0 | 0 | 0 | 330 |
| Candidatus Brocadaceae | 457 | 0 | 0 | 249 | 236 |
| Segniliparaceae | 423 | 0 | 0 | 132 | 332 |
| Schistosomatidae | 416 | 0 | 0 | 296 | 524 |
| Desulfobulbaceae | 415 | 0 | 0 | 126 | 406 |
| Erythrobacteraceae | 400 | 0 | 0 | 0 | 382 |
| Piscirickettsia group | 399 | 0 | 0 | 102 | 248 |
| Pelobacteraceae | 396 | 0 | 0 | 249 | 588 |
| Solibacteraceae | 390 | 0 | 0 | 0 | 286 |
| Dasyuridae | 372 | 0 | 0 | 400 | 358 |
| Helibacteriaceae | 371 | 0 | 0 | 198 | 502 |
| Acaridae | 341 | 0 | 0 | 111 | 217 |
| Rhodothermaceae | 334 | 0 | 0 | 0 | 722 |
| Iridoviridae | 333 | 0 | 0 | 0 | 0 |
| Cyclobacteriaceae | 328 | 0 | 0 | 184 | 264 |
| Desulfarculaceae | 317 | 0 | 0 | 149 | 440 |
| Trueperaceae | 309 | 0 | 0 | 104 | 200 |
| Dictyoglomaceae | 298 | 0 | 0 | 147 | 166 |
| Phycisphaeraceae | 296 | 0 | 0 | 135 | 370 |
| Desulfobacteraceae | 282 | 0 | 0 | 0 | 343 |
| Vibrionaceae | 277 | 0 | 0 | 0 | 400 |
| Gallionella group | 275 | 0 | 0 | 0 | 311 |
| Hydrogenothermaceae | 275 | 0 | 0 | 118 | 156 |
| Ignavibacteriaceae | 268 | 0 | 0 | 0 | 155 |
| Methylococcaceae | 256 | 0 | 0 | 0 | 296 |
| Shewanellaceae | 252 | 0 | 0 | 102 | 161 |
| Nostocaceae | 240 | 0 | 0 | 0 | 113 |
| Nitrosomonadaceae | 229 | 0 | 0 | 104 | 271 |
| Hydrogenophilaceae | 227 | 0 | 0 | 0 | 209 |
| Orbaceae | 223 | 0 | 0 | 158 | 189 |
| Acidithermaceae | 218 | 0 | 0 | 0 | 188 |
| Caviidae | 218 | 0 | 0 | 0 | 0 |
| Parvularculaceae | 218 | 0 | 0 | 0 | 137 |
| Planctomycetaceae | 217 | 0 | 0 | 111 | 206 |
| Tetrahymenidae | 214 | 0 | 0 | 0 | 0 |
| Natronaerobiaceae | 212 | 0 | 0 | 132 | 209 |
| Nitrospiraceae | 211 | 0 | 0 | 0 | 136 |
| African mole-rats | 207 | 0 | 0 | 104 | 158 |
| Caldilineaceae | 207 | 0 | 0 | 100 | 224 |
| Deferribacteraceae | 207 | 0 | 0 | 279 | 268 |
| Syntrophaceae | 205 | 0 | 0 | 115 | 216 |
| Brevibacteriaceae | 200 | 0 | 0 | 0 | 0 |
| Gemmantimonadaceae | 199 | 0 | 0 | 0 | 139 |
| Desulfomicrobiaceae | 194 | 0 | 0 | 103 | 219 |
| Strongylocentrotidae | 186 | 0 | 0 | 0 | 0 |
| Thermoanaerobacterales Family IV. Ir | 186 | 0 | 0 | 109 | 222 |

**Supplemental table S5: Summary of classification to the family level for all samples**
|  |  |  |  |  |  |
| --- | --- | --- | --- | --- | --- |
| Fabaceae | 184 | 0 | 0 | 0 | 389 |
| Chitinophagaceae | 178 | 0 | 0 | 0 | 239 |
| Thermodesulfobacteriaceae | 170 | 0 | 0 | 154 | 203 |
| Desulfurobacteriaceae | 168 | 0 | 0 | 0 | 104 |
| Sulfuricellaceae | 163 | 0 | 0 | 0 | 0 |
| Legionellaceae | 160 | 0 | 0 | 0 | 0 |
| Colwelliaceae | 159 | 0 | 0 | 0 | 144 |
| Entomoplasma group | 159 | 0 | 0 | 0 | 152 |
| Syntrophomonadaceae | 158 | 0 | 0 | 0 | 123 |
| Camelidae | 154 | 0 | 0 | 0 | 209 |
| Debaryomycetaceae | 152 | 0 | 0 | 174 | 0 |
| Cryomorphaceae | 151 | 0 | 0 | 113 | 273 |
| Bdellovibrionaceae | 145 | 0 | 0 | 0 | 142 |
| Ferrimonadaceae | 145 | 0 | 0 | 0 | 105 |
| Saprospiraceae | 144 | 0 | 0 | 0 | 166 |
| Roseiflexaceae | 140 | 0 | 0 | 0 | 324 |
| Chrysiogenaceae | 139 | 0 | 0 | 0 | 196 |
| Tetraodontidae | 139 | 0 | 0 | 0 | 0 |
| Chloroflexaceae | 133 | 0 | 0 | 0 | 0 |
| Draconibacteriaceae | 130 | 0 | 0 | 0 | 117 |
| Cneoraceae | 127 | 0 | 0 | 200 | 300 |
| Culicidae | 124 | 0 | 0 | 0 | 131 |
| Flammeovirgaceae | 117 | 0 | 0 | 0 | 0 |
| Hypocreaceae | 113 | 0 | 0 | 0 | 0 |
| Leeaceae | 112 | 0 | 0 | 216 | 218 |
| Rivulariaceae | 111 | 0 | 0 | 0 | 0 |
| Arthrodermataceae | 110 | 0 | 0 | 0 | 0 |
| Desulfurellaceae | 107 | 0 | 0 | 0 | 117 |
| Hahellaceae | 107 | 0 | 0 | 0 | 107 |
| Acidithiobacillaceae | 103 | 0 | 0 | 0 | 247 |
| Noelaerhabdaceae | 103 | 0 | 0 | 512 | 379 |
| Pinaceae | 103 | 0 | 0 | 0 | 0 |
| Sphaeriaceae | 103 | 0 | 0 | 0 | 0 |
| Alligatoridae | 0 | 0 | 0 | 0 | 253 |
| Balaenopteridae | 0 | 0 | 0 | 0 | 754 |
| Chaetomiaceae | 0 | 0 | 0 | 241 | 274 |
| Costariaceae | 0 | 0 | 0 | 130 | 0 |
| Dipodascaceae | 0 | 0 | 0 | 220 | 0 |
| Drosophilidae | 0 | 0 | 0 | 0 | 178 |
| Fibrobacteraceae | 0 | 0 | 0 | 0 | 140 |
| Francisella group | 0 | 0 | 0 | 0 | 146 |
| Hydriidae | 0 | 0 | 0 | 0 | 181 |
| Lagriidae | 0 | 0 | 0 | 0 | 109 |
| Loliginidae | 0 | 0 | 0 | 0 | 171 |
| Magnetococcaceae | 0 | 0 | 0 | 0 | 102 |
| Mamiellaceae | 0 | 0 | 0 | 0 | 161 |
| Mycosphaerellaceae | 0 | 0 | 0 | 349 | 1,122 |
| Mycosyringaceae | 0 | 0 | 0 | 0 | 228 |
| Onchocercidae | 0 | 0 | 0 | 0 | 119 |
| Pseudoalteromonadaceae | 0 | 0 | 0 | 127 | 231 |
| Retroviridae | 0 | 0 | 0 | 296 | 174 |
| Rhabditidae | 0 | 0 | 0 | 126 | 0 |
| Sarcocystidae | 0 | 0 | 0 | 261 | 140 |
| Syntrophobacteraceae | 0 | 0 | 0 | 0 | 119 |
| Thermodesulfobiaceae | 0 | 0 | 0 | 0 | 136 |
|  | 26,519,857 | 386,315 | 342,077 | 14,241,186 | 25,157,407 |

